# Multi-region two-photon calcium imaging in freely moving animals

**DOI:** 10.64898/2026.10.27.760324

**Authors:** Embla Kjellevold-Skallerud, Shouwen Ma, Julija Arbačiauskaitė, Rajat Saxena, Antonio Lorca Camara, Klaus Jøran Jenssen, Jonathan R. Whitlock, Manfred Gahr, Weijian Zong

**Author notes:** These authors contributed equally (Embla Kjellevold-Skallerud, Shouwen Ma). Correspondence (W.Z.).

## Abstract

Complex behaviors emerge from the coordinated activity of neural circuits distributed across widely separated brain regions. Existing optical approaches for monitoring neural activity lack the combined capability to record from anatomically distributed regions with high spatiotemporal resolution in freely moving animals. Here we introduce MINI2P-II, a 1.1-g 2P miniscope that achieves large-scale calcium imaging with high spatial resolution, deep imaging depth and dual-color imaging capability. Together with a dual-scope imaging system and a novel implantable periscope lens assembly, MINI2P-II enables simultaneous 2P imaging in widely separated or anatomically adjacent brain regions. In mice, we demonstrate inter-hemisphere coordination in bilateral motor cortex during climbing, egocentric boundary coding in simultaneous retrosplenial and posterior parietal cortex recordings, and in zebra finches, we showed large-scale behavior-dependent activity in both juvenile and adult individuals. MINI2P-II opens the door to multi-region 2P imaging across species and developmental stages in freely moving animals.

## Introduction

A fundamental question in neuroscience is how interactions between neural circuits give rise to complex behaviors (*1*). A major challenge in addressing this question is the development of tools that can monitor large-scale neural activity from distributed brain areas across different behavioral states. An ideal solution is a miniaturized recording instrument that provides high cell yield, fine spatial resolution and fast acquisition speed, while remaining sufficiently lightweight to be carried by freely moving animals without disrupting naturalistic behavior. Animals have been shown to maintain natural behavior with head-mounted devices less than 10% of their body weight (*2, 3*). Following this guideline, minimizing weight allows for a greater number of devices that can be mounted on the head to access multiple brain regions simultaneously.

Multi-channel electrophysiological probes are light and enable simultaneous recordings across diverse brain structures (*4, 5*). However, recording probes can face problems of glial encapsulation, signal degradation, and losing track of the same neurons over days to weeks, making long-term studies difficult. While calcium imaging does not match the temporal resolution of electrophysiological probes, it allows for optically measuring neuronal morphology and intracellular calcium transients linked to action potentials (*6*). One-photon (1P) miniscopes (*7, 8*) have achieved remarkable miniaturization (e.g. weighing less than 1 gram), which enabled multi-region imaging in freely moving mice (*9, 10*). However, 1P imaging suffers from background contamination, limited penetration and photobleaching, making it less suitable for dense labeling and long-term recording (*11*). Moreover, ultralight (sub-1-g) designs typically rely on low-quality optics, which further reduce spatial resolution and cell yield.

By contrast, miniature two-photon (2P) microscopes (i.e. 2P miniscopes) (*3, 12–15*) promise to overcome many of these limitations. 2P miniscopes rely on near-infrared excitation and nonlinear absorption, which provide excellent optical sectioning, reduced background and lower photobleaching (*16, 17*). Recent developments of 2P miniscopes have led to improved optical performance, making them rapidly approach the spatiotemporal-resolution (*15, 18*), cell yield (*3, 19*) and multi-color imaging capability (*20*) of benchtop 2P systems. However, the weight of high-performance 2P miniscopes has not been able to break below 2 grams (*21*). As a result, experiments are typically restricted to a single device mounted on adult mice weighing 20-30 g, with smaller species or early developmental stages being off limits. Attempts to further decrease the weight of 2P miniscopes are emerging but, like ultralight 1P systems, these systems sacrifice spatial resolution and cell yield (*22, 23*). Such trade-offs between imaging performance and weight reduction suggested a fundamental limitation, which is unlikely to be broken via current paths of technical development. Instead, new design concepts are needed.

Here we overcome this limitation by introducing MINI2P-II, a novel 2P miniscope weighing only 1.1 g. Based on a new optical framework, MINI2P-II achieved such extreme miniaturization with a similar field-of-view (FOV) size, speed, Z-scanning capability, and improved spatial resolution and detection efficiency compared to our previous MINI2P system (*3*).

Beyond weight reduction, we developed ready-to-use solutions for simultaneous, multi-region imaging both contra- and ipsilaterally in freely moving mice. These include complementary approaches for recording widely distributed regions, along with a new implantable optical element—the GRIN periscope assembly (GPA)—that targets neighbouring regions as close as 1 mm. We provide three key demonstrations for MINI2P-II, showing its wide applicability. First, we simultaneously record bilaterally from primary motor cortex (M1) during naturalistic behaviors (climbing, eating, running, etc.), revealing behavior-dependent population dynamics across hemispheres. Second, we recorded the ipsilateral retrosplenial cortex (RSC) and posterior parietal cortex (PPC) simultaneously, enabling direct comparisons of egocentric boundary tuning across regions, both within animals and within the same session. Finally, to showcase applications in smaller species and early developmental stages, we performed high-quality calcium imaging in freely moving zebra finches: a 130-day-old adult male (16 g) during courtship singing and a 44-day-old juvenile (14 g) during social interactions.

Together, these results demonstrate that the MINI2P-II enables multi-region 2P calcium imaging in freely moving animals and expands the behavioral paradigms, species, and developmental stages accessible to 2P imaging. The system is fully open source and ready for broad adoption by the neuroscience community.

## Results

### New design concepts to break the weight bottleneck

Across different 2P miniscope technical paths, the approach based on micro-electro-mechanical-system (MEMS) scanners (i.e. fulfilling 2P scanning via a MEMS mirror) provides state-of-the-art imaging performance and is under the most active development (*21*). We therefore used this approach as the skeleton for the MINI2P-II design. Three major factors determine the minimum weight and size of a MEMS-based 2P miniscope: the number of optical elements, the volume of supporting structures (the “scope body”) that maintain the distance between the optical elements required for image formation, and the mass of each element. We systematically optimized MINI2P-II in each aspect.

To reduce the number of optical elements while retaining essential imaging functions, we asked whether certain elements could be shared. Specifically, could a single element replace several elements that require similar optical parameters? MEMS-based 2P miniscopes (*3, 15, 18–20*) need a fiber collimator (which collimates the diverging laser beam from the fiber) and a scan lens (which converts angular deflections of the laser beam by the MEMS scanner into lateral positional shifts in the imaging plane). These two elements typically require nearly identical parameters: effective focal length (EFL), numerical aperture (NA), and near-infrared operating wavelength. Therefore, in the new miniscope we used only a telecentric f-theta lens (TFL) as both the fiber collimator and the scanning lens (**Figs. 1A-i, 1B)**.

**Figure 1.**
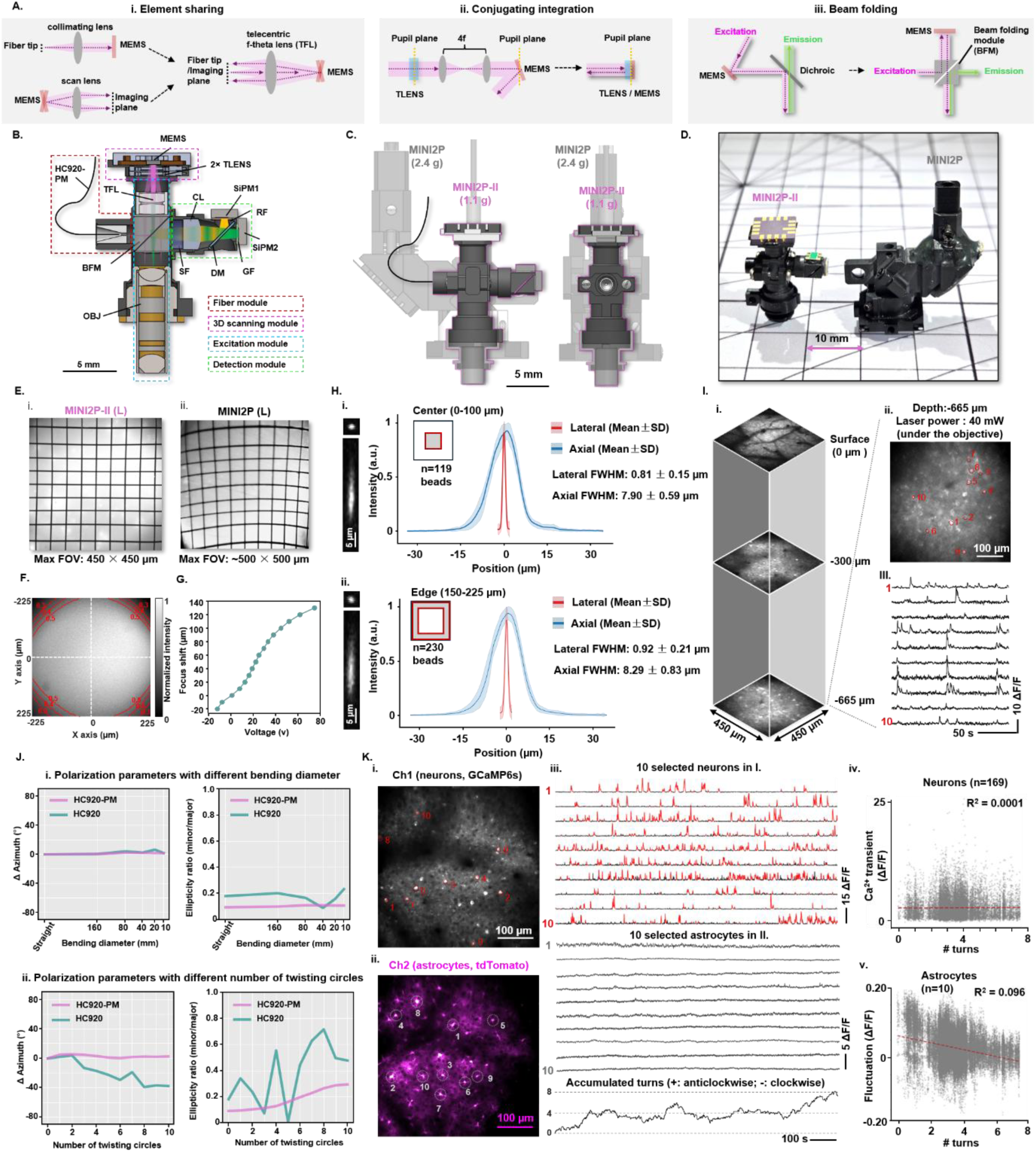
MINI2P-II achieving low weight, small size, and excellent optical performance. A) Three core design concepts enabling weight and size reduction in MINI2P-II: (i) element sharing, (ii) conjugated integration of scanning optics, and (iii) beam folding. B) Optical schematic of MINI2P-II showing key components. HC920-PM, polarization-maintaining hollow-core fiber; OBJ, objective; SF, short-pass filter; CL, collection lens; DM, dichroic mirror; GF, green emission filter (500–550 nm); RF, red emission filter (600–650 nm); SiPM, silicon photomultiplier; C) Overlay of MINI2P and MINI2P-II drawings, illustrating the reduction in overall size and mass (MINI2P ∼2.4 g; MINI2P-II ∼1.1 g). D) Photograph of MINI2P-II and MINI2P side-by-side. Note the smaller footprint of MINI2P-II (mounting area on skull) relative to MINI2P. E) Maximum field of view (FOV) quantified with a 50-µm grid sample for MINI2P-II (i) and MINI2P (ii). Note the reduced image distortion (top–bottom grid size difference) with MINI2P-II(L). F) FOV homogeneity measured on a uniform fluorescent slide at the maximum imaging area (450 × 450 µm). Red contours mark regions at 50%, 40%, and 30% of peak intensity. G) Axial (Z) scanning range of MINI2P-II using two tunable lenses integrated within the 3D scanning module. H) Spatial resolution at the center (i) and edge (ii) of the FOV. Left, example bead images with XY and YZ cross-sections. Right, normalized averaged profiles (mean ± SD) for lateral (red) and axial (blue) point-spread functions (PSFs) from N = 119 beads at center (i) or 230 beads at edge (ii), lateral and axial FWHM of Gaussian fitting curves are extracted. Note the minimal PSF degradation toward the FOV edge. I) Demonstration of deep-tissue functional calcium imaging with MINI2P-II. (i) Mean-intensity projections at 0 µm (surface), −300 µm, and −665 µm in visual cortex (VC) of a GCaMP6s mouse. (ii) Averaged image at -665 µm. (iii) Ten example neuronal calcium traces corresponding to ROIs in (ii). Note the high SNR at -665 µm using only 40 mW laser power (measured under the objective). J) Polarization stability of HC920-PM versus HC920 fibers (length 2.3 m) under bending and twisting. (i) Changes in azimuth (left) and ellipticity ratio (right) when bending from straight to a 10-mm diameter loop. (ii) Changes in azimuth (left) and ellipticity ratio (right) when twisting from 0 to 10 full turns. Note that HC920-PM preserves polarization substantially better than HC920 (smaller azimuth shifts and reduced ellipticity changes) under severe deformation. K) Stability of dual-color MINI2P-II imaging with HC920-PM in a freely moving mouse over 18 min (V1, depth 200 µm, also watch Video S1). (i) Average image of GCaMP6s-expressing neurons (green channel). (ii) Average image of tdTomato-labeled astrocytes (red channel). (iii) Top: ten example neuronal ΔF/F traces (ROIs as in k-I, Red parts of traces: suprathreshold calcium transients). Middle: ten example astrocyte intensity traces (ROIs as in k-II). Bottom: accumulated cable-twist angle over time (positive = clockwise; negative = counter-clockwise). (iv–v) Linear-regression analysis between cable twisting and (iv) neuronal calcium-transient amplitude (N = 169 neurons, ΔF/F in all frames falling in red periods in III-top) or (v) astrocyte mean intensity (N = 10 astrocytes, ΔF/F in all frames). Gray dots: single frame; red dashed line: linear fitting. Low R² indicates no correlation, supporting stable imaging despite cable twisting.

To shorten distances between elements while preserving their required optical relationships (such as conjugation and magnification), we identified two components that should be at conjugated positions, i.e., the MEMS scanner and the Tlenses(*3*), and integrated them into a single 3D scanning module (**Figs. 1A-ii, 1**, and Step 1 in **S1H**). This integration ensures that incident light passes paraxially through the Tlenses and hits the MEMS mirror at normal incidence, both increasing the usable aperture of the MEMS mirror and eliminating the scanning distortion observed in earlier 2P miniscopes (**Fig. 1E**) (*3, 15*). It also reduces the number of Tlenses required for large-range Z-scanning by allowing the beam to pass twice through the same Tlenses (e.g. 150-micron with 2 Tlenses, **Fig. 1F**).

Next, we introduced a beam folding module (BFM, **Fig. S1A**), which allows different light paths to occupy the same physical space while traveling in different directions depending on polarization state and wavelength (**Fig. 1A-iii**). The BFM makes the “element-sharing” and “conjugated-integration” concepts feasible (**Figs. 1B, S1B**, and **S1C**).

Finally, beyond reducing the number of elements and minimizing spacing, we decreased component weight. We customized a smaller MEMS package from Mirrorcle (CA, USA) that weighs less than half of the previous version (**Fig. S1D-i**) (*3*), and we replaced stainless-steel housings for the objective and TFL with PEEK-CF30 (a lightweight carbon-fiber-reinforced polymer; **Fig. S1D-ii**).

With these improvements, MINI2P-II achieved a total weight of only 1.1 g including all components for two-color imaging, except the fiber module, the electronic cables and the baseplate (**Figs.1C**, **S1E**). Its footprint, i.e., the space on the skull required for mounting, was reduced to a diameter of 5 mm. Both weight and footprint are less than half that of the original MINI2P (weight: 2.4 g, footprint: 11 × 11 mm^2^, **Figs. 1C** and **1D**).

### MINI2P-II achieves excellent optical performance, deep imaging depth, and high modularity for convenient function upgrade

MINI2P-II reached a maximum FOV of 450 × 450 µm² using the large-angle-version MEMS (MEMS-L) as used in MINI2P (**Fig. 1E**), with high homogeneity (82.6% of the FOV maintains ≥50% of peak intensity, 95.5% ≥ 30%, Fig.**1F**), a Z-scanning range of 150 µm (**Fig. 1G**), and spatial resolution of <1 µm laterally and ∼8 µm axially (**Fig. 1H**, mean ± SD; **i**, center: 0.81 ± 0.15 µm XY, 7.90 ± 0.59 µm Z; **ii**, edges: 0.92 ± 0.21 µm XY, 8.29 ± 0.83 µm Z). The improved axial resolution relative to the MINI2P (8 µm vs. 12-15 µm) (*3*) results from a shorter focal length of the scan lens (so, higher excitation NA), and an optimized TFL design that reduces optical aberrations.

Using onboard silicon photomultipliers (SiPMs; **Fig. 1B**) also provides higher signal detection efficiency(*19, 24*) than fiber-bundle-based detection, which is still commonly used in 2P miniscopes (*3, 15, 18, 20*). And it allows MINI2P-II to maintain a high signal-to-noise ratio (SNR) even when imaging deep brain tissue. We verified deep-tissue functional imaging by recording in visual cortex through a glass window in a head-fixed CaMKII-tTA/tetO-GCaMP6s mouse (**Fig. 1I**), observing high-SNR Ca²⁺ transients at 665 µm depth using only 40 mW laser power under the objective (**Fig. 1I**).

The MINI2P-II was also designed for highly functional modularity. The microscope can be divided into four main functional modules: the fiber module, 3D-scanning module, excitation module, and detection module (**Fig. 1B**). The mounting interfaces for these modules are positioned orthogonally to each other and standardized (M1.2-6 standard, see Methods for details), which allows easy replacement of individual modules and integration of additional functional modules in between (**Fig. S1F**). Several key components, including fibers, MEMS scanners, and objectives have multiple commercial options (**Supplementary Documents Part 3: Part list**). The threading and mount-to-image-plane distance of objective are compatible with MINI2P and all miniature objectives from the Thorlabs catalog.

### A new polarization-maintaining hollow-core fiber enables stable recording in freely moving mice

Using polarization-dependent optics in freely moving conditions requires that the laser polarization output from the fiber remains stable under bending and twisting, conditions that frequently occur during unrestrained animal behavior. The original hollow-core fiber (HC920)(*15*) does not maintain polarization, so was unlikely to satisfy this requirement (**Fig. S1G-i**). To overcome this, we customized a new polarization-maintaining hollow-core fiber (HC920-PM) with higher birefringence (≥ 2.8 × 10⁻⁴ in HC920-PM vs. < 0.9 × 10⁻⁴ in HC920; **Fig. S1G-i**) from NKT (now Hamamatsu Photonics, Denmark).

To evaluate the polarization-maintaining performance of HC920-PM compared with HC920, we first performed controlled bending and twisting experiments. We bent 2.5-meter fiber segments to defined diameters (**Fig. S1G-ii**) and twisted them by specific numbers of full turns (**Fig. S1G-iii**), while continuously monitoring the polarization state of the laser output (**Fig. S1G-iv**, **v**, center wavelength: 920 nm). Two parameters were extracted: Δazimuth (the change in the orientation of the linear polarization axis relative to a reference; **Fig. 1J**) and the ellipticity ratio (0 indicates perfectly linear polarization; 1 indicates perfectly circular). When bent down to a diameter of 10 mm (**Fig. S1G-ii**), both fibers maintained stable polarization (**Figs. 1J-i**, **S1G-iv**), but HC920-PM showed smaller azimuth and ellipticity changes than HC920. When twisted up to 10 full rotations, representing the most severe realistic torsion in freely moving recordings (**Fig. S1G-iii**), the HC920-PM fiber again preserved polarization far better than HC920 (**Figs. 1J-ii**, **S1G-v**), with small and incremental azimuth shift and ellipticity change. In contrast, the changes for HC920 were larger and unpredictable.

To test whether MINI2P-II combined with HC920-PM maintains stable imaging in actual freely moving conditions, we performed dual-color MINI2P-II imaging in primary visual cortex (V1) of a female mouse freely foraging in an 80-cm arena (**Figs. 1K** and **Video S1**). Neuronal Ca²⁺ activity (GCaMP6s, green channel; **Fig. 1K-i**) and astrocyte tdTomato intensity (red channel; **Fig. 1K-ii**) remained stable throughout the 18-min recording, even during substantial movement of the animal (**Fig. 1K-iii; Video S1**). We quantified the relationship between fiber twist angle (accumulated turns over time) and signal stability using linear regression (**Fig. 1K-iv** and **v)**. No significant association was found between accumulated twist angle and neuronal calcium-transient amplitude (ΔF/F; *R²* = 0.0001, **Fig. 1K-iv**) or between twist angle and astrocyte fluorescence intensity (*R²* = 0.096, **Fig. 1K-v**), indicating that HC920-PM maintains polarization in real behavioral conditions and enables stable, high-quality dual-color imaging.

In summary, we developed a new generation of 2P miniscope—MINI2P-II—that combines low weight, small size, and excellent optical performance. MINI2P-II surpasses the MINI2P in spatial resolution and detection efficiency. It can be easily replicated in biology labs following the assembly protocol (**Fig. S1H**) and complete parts list (**Supplementary Document Part 3: parts list**). It is modular and scalable, providing tool developers with a standardized platform for future functional extensions.

### Using two or more MINI2P-II miniscopes simultaneously

Building on MINI2P-II’s ultralight design, we developed an open-source dual-scope system for simultaneous recordings from multiple brain regions in freely moving mice (**Figs. S2A** and **S2B**), including a compact beam-splitting/dual-fiber-coupling module that delivers laser light to two miniscopes from a single laser source (**Fig. S2C**). The system operates natively with ScanImage software, controlling two miniscopes with two-color detection in parallel, or up to four miniscopes in single-channel mode. For labs with limited engineering capacity, commercial laser systems (e.g., ALCOR MULTIBEAM, SparkLaser, France) are available, offering one independent laser power control per fiber output and up to four fiber outputs that are compatible with MINI2P-II, enabling simultaneous imaging with as many as four miniscopes (**Fig. S2G**).

Simultaneous imaging across widely separated regions typically requires separate cranial windows (**Figs. S2H** and **S2I**). To accommodate this, we designed a multi-miniscope mounting station to align each miniscope independently and in parallel on the animal’s head (**Fig. S2D**). Using this station, each miniscope is aligned to its corresponding headbar, and thus to its window, without interference, maximizing regional flexibility and avoiding the need to measure inter-window angles (**Fig. S2H and S2I**, also see Methods for details).

### Carrying two MINI2P-II miniscopes with connection cables does not impair naturalistic behavior

Our previous work showed that maintaining naturalistic movement in adult mice during 2P imaging required both ≤3 g head-mounted mass and thin, flexible cables (*3*). Even though having two mounted MINI2P-II devices satisfies this weight criterion, the stiffness and weight of the cables can still constrain behavior (3). Indeed, in our pilot experiments, using two separate cables (9 wires for each miniscope; 18 total) reduced the animals’ locomotion. We solved this issue by designing a new cable solution that shares several electronic signals between two miniscopes, decreasing the total number of wires to 12 (a net increase of 3 wires from one miniscope), while maintaining functions that need to be run individually for each miniscope (**Figs. 2A, S2E** and **S2F,** also see Method section “Electronic cable assembly” for details).

**Figure 2.**
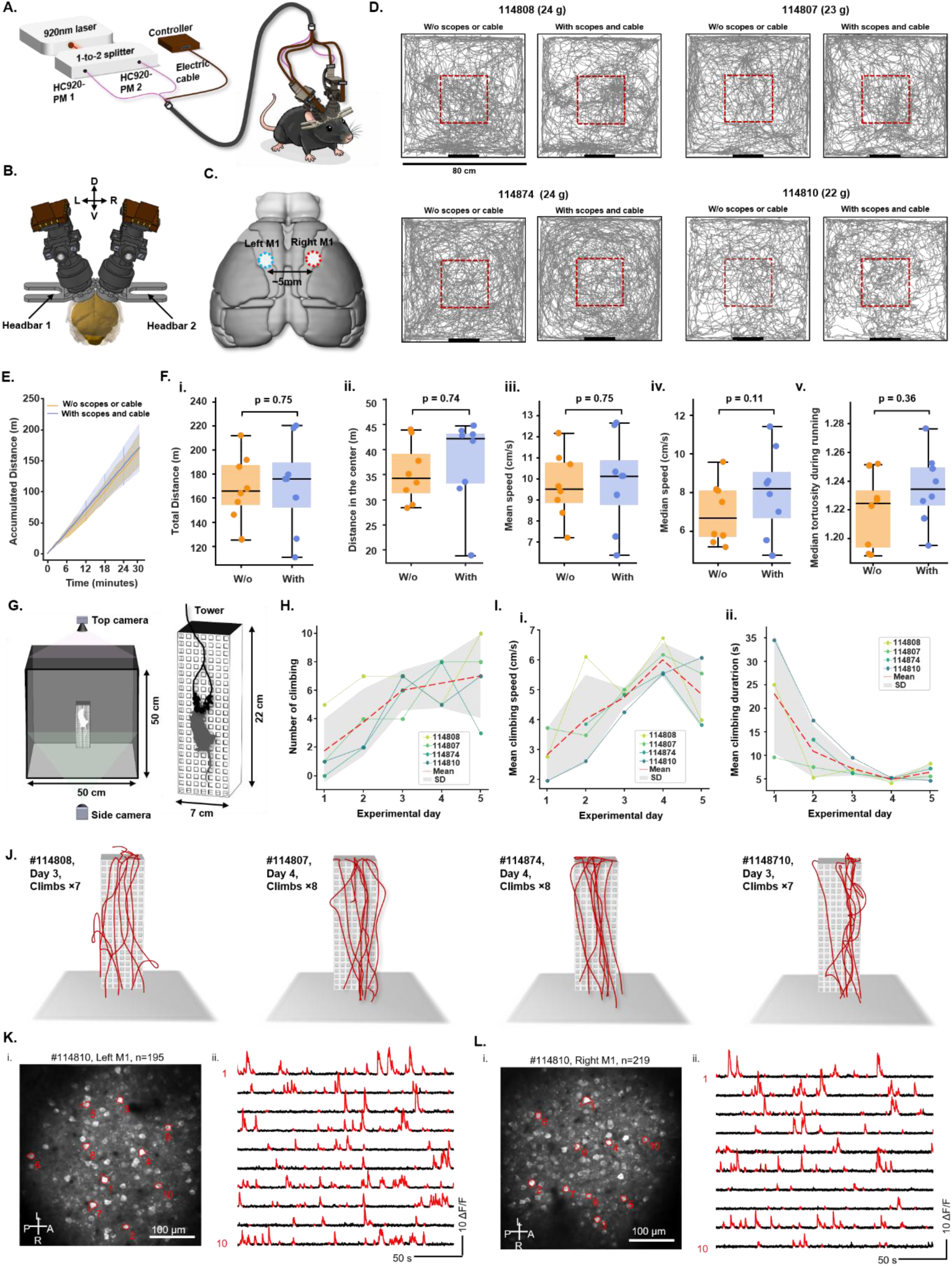
Dual-MINI2P-II system enables simultaneous two-site imaging without restricting natural movement. A) Simplified system schematic for simultaneous recordings from two brain regions in freely moving mice. A single 920-nm laser feeds a compact beam-splitting/dual-fiber coupling module that launches light into two HC920-PM fibers. Two MINI2P-II miniscopes are mounted on the head; a thin shared electrical cable delivers MEMS and SiPM control signals. B) Illustration of mounting two miniscopes on left and right hemispheres, each aligned to its own headbar for independent positioning. C) Cortical targets from a dorsal view: miniscope 1 on left primary motor cortex (M1, blue dashed circle) and miniscope 2 on right M1 (red dashed circle). D) Representative trajectories from four mice freely exploring an 80 × 80 cm open field. Each mouse performed two 29-min sessions: “W/o scopes or cable” (no miniscope/cable) and “With scopes and cable” (two miniscopes plus connection cable). Thick black line, cue card. Mouse ID and body weight are indicated. Red dashed boxes show 30 × 30 cm^2^ squares used in **F**. E) Accumulated distance over 29 min. Lines, mean across four mice; vertical bars at six time points, SD; shaded band, SD across all time points. Note the similarity between conditions. F) Box plots comparing locomotion metrics between conditions (29-min sessions). i, total distance; ii, distance in the central 30 × 30 cm square; iii, mean speed; iv, median speed; v, median tortuosity during running (speed > 2.5 cm/s). Horizontal line, median; box, interquartile range (IQR); whiskers, 1.5× IQR; colored dots, individual mice. Statistics: Shapiro–Wilk test for normality; paired t-test when normal distribution was assumed and Wilcoxon signed-rank test otherwise. P values shown on plots. G) Ladder-climbing assay. Mice were trained to climb up and then climb/jump down from a 3D-printed PLA tower (22-cm height; 7 × 7 cm top). One vertical face (22 × 7 cm) contains square holes (6 × 6 mm²) spaced 4 mm apart (ladder). The arena (50 × 50 cm) has a black rubber floor and 50-cm-high black acrylic walls. Two near-infrared cameras recorded behavior: camera1, top view; camera2, front view through a wall aperture aligned to the ladder. H) Number of successful climbs (reaching the top of the platform) across training days. I) Performance with two MINI2P-II devices stabilizes across training. (i) Mean climbing speed increases and stabilizes across trials; (ii) climb duration to the top decreases across days. J) Example head trajectories of four mice during climbing on day 3 (mouse #114808 and #114810) and day 4 (mouse #114807 and #114874) of the experiment. K) Left M1 imaging example during ladder climbing task. (i) Maximum-intensity projection from a single field of view (FOV) in left primary motor cortex (M1); 195 active neurons were identified by manually examining the Suite2p output. (ii) Ten example ΔF/F traces from neurons marked in **K-i** (Red parts of traces: suprathreshold calcium transients). L) Right M1 example (same session as k). (i) Maximum-intensity projection from right M1; 219 active neurons identified. (ii) Ten example ΔF/F traces from neurons marked in **L-I**.

To test whether head-mounting of two MINI2P-II miniscopes and new cables compromise normal locomotor behavior of freely moving animals, we used four adult transgenic mice (CaMKII-GCaMP6s; 22–24 g, all male) implanted with two miniscopes, one in each hemisphere, following the surgery protocol described above. We specifically targeted the primary motor cortex, (M1; **Figs. 2B, 2C**; coordinates: AP +1.0 mm, ML 2.5 mm left and right of midline) due to its surgical accessibility and role in driving behavior. Each mouse was allowed to forage freely for scattered food crumbs during 30-min sessions in an 80-cm square open field, either with both miniscopes and cables attached (“With” sessions) or without any miniscope or tether (“W/o” sessions; **Figs. 2D** to **2F**).

All animals showed robust running behavior in both conditions (**Fig. 2D**) and continued to explore and forage throughout the sessions (**Fig. 2E**). Total distance traveled, distance in the center zone, mean and median speed did not differ significantly between tethered and untethered sessions (Shapiro–Wilk test for normality; paired t-tests or Wilcoxon signed-rank tests as appropriate. Total distance: p = 0.75, **Fig. 2F-i**; center-zone distance: p = 0.74, **Fig. 2F-ii**; mean speed: p = 0.75, **Fig. 2F-iii**; median speed: p = 0.11, **Fig. 2F-iv**). These values were consistent with previous behavioral measurements reported for the MINI2P (*3*). Trajectory tortuosity, a metric of path complexity (Methods) also showed no difference between conditions (paired t-test: p = 0.36; **Fig. 2F-v**).

To further examine whether behavior was affected during tasks requiring more complex movement sequences, we trained the same four mice to climb a ladder vertically — a task we developed previously to assess the impact of head-mounted miniscopes on vigorous behavior (*3*). In each experiment, the mouse was placed in a 50 cm square box containing a raised platform (7 × 7 cm² base; 22 cm height; tower in **Fig. 2G** right) positioned near the center of one wall (**Fig. 2G** left). Only one face of the tower was fitted with a ladder (a grid of 7 × 7 mm² square apertures with 3 mm spacing), facing toward the center of the box. A side camera was placed outside the box to monitor climbing, and an additional top camera was mounted above the box to track locomotion on the floor. In the task, mice climb vertically up a ladder to reach a small reward on the top of the tower, then climb or jump down to begin the next bout (**Video S2**). After only 3 days of training, all 4 mice completed at least 5 successful climbs (reaching the top of the platform) per session with two miniscopes mounted (∼30 min; **Figs. 2H** and **2J**). Average climbing speed increased (**Fig. 2I-i**), and the time required to reach the top decreased and stabilized across days (**Fig. 2I-ii**).

Together, these observations suggest that mounting two MINI2P-II miniscopes with optimized connection cables does not disrupt naturalistic behaviors we examined. The animals maintained normal locomotion, exploration and could fulfill difficult vertical climbing tasks.

### High-quality simultaneous bihemispheric calcium imaging during vigorous behavior

After confirming that carrying two MINI2P-II miniscopes did not affect exploratory behavior, we asked whether the dual-scope imaging system delivered high imaging quality and stability from both miniscopes during vigorous and varied motor behaviors. We addressed this question using the ladder climbing sessions described above, recording neural activity from large populations of neurons using MINI2P-II miniscopes implanted in homotopic regions of M1 in the left and right hemispheres (see **Supplementary Document, Part 2**). We used transgenic GCaMP6s mice to ensure broad expression in excitatory neurons throughout the brain, and to minimize regional variation in indicator expression and baseline activity levels.

In an example session (**Figs. 2K**, **2L**; **Video S2**) we recorded 195 neurons from left M1 (**Fig. 2K**) and 219 neurons from right M1 (**Fig. 2L**), each in a single plane at layer II. Neurons exhibited large numbers of calcium transients with high ΔF/F amplitudes (example neurons shown in **Fig. 2K-ii** and **Fig. 2L-ii**). Across all four mice and 16 recordings, we consistently observed stable numbers of detected cells (**Fig. S2J-i**), high SNR (**Fig. S2J-ii**) and low levels of motion-related displacement comparable to those typically observed in MINI2P imaging (*3*) (**Fig. S2J-iii and iv**).

No significant differences were observed for these factors across left and right hemispheres (Mann-Whitney U test, n = 16, U = 119.0, p = 0.75 for cell numbers, **Fig.S2J-i**, U = 128.0, p = 1.00 for median SNR, **Fig.S2J-ii,** U = 123.0, p = 0.78 for FOV shift in x axis, **Fig.S2J-iii**, and U = 94.0, p = 0.21 for FOV shift in y axis, **Fig.S2J-iv**).

Together, these results demonstrated that our dual-scope system enables large-scale, simultaneous two-region calcium imaging during vigorous naturalistic behavior. We now show over the following 5 sections how this system, combined with the novel vertical climbing paradigm, opens new experimental windows into population coding in the motor cortex and inter-hemispheric coordination across cortices.

### 2Using two MINI2P-II to study inter-hemispheric coordination during reward-motivated climbing

#### i. Population 2P imaging in bihemispheric forelimb motor cortex during a ladder climbing task

Previous studies using relatively simple and controlled motor tasks, such as lever pressing (*25*) and head-fixed treadmill locomotion (*26*) have shown that limb-specific movements engage bilateral M1 coordination. However, whether the structure of this bilateral engagement changes with behavioral state or movement complexity during naturalistic behavior remains much less clear. We shed new light on this by combining reward-motivated ladder climbing, consisting of well-defined behavioral phases and quantifiable behavioral states (**Figs. 3A** and **S3A**), with high-quality, projection-defined bilateral imaging in M1. Using this approach, we uncovered patterns in inter-hemispheric coordination that would be impossible to measure with unilateral recordings or head-fixed paradigms.

**Figure 3.**
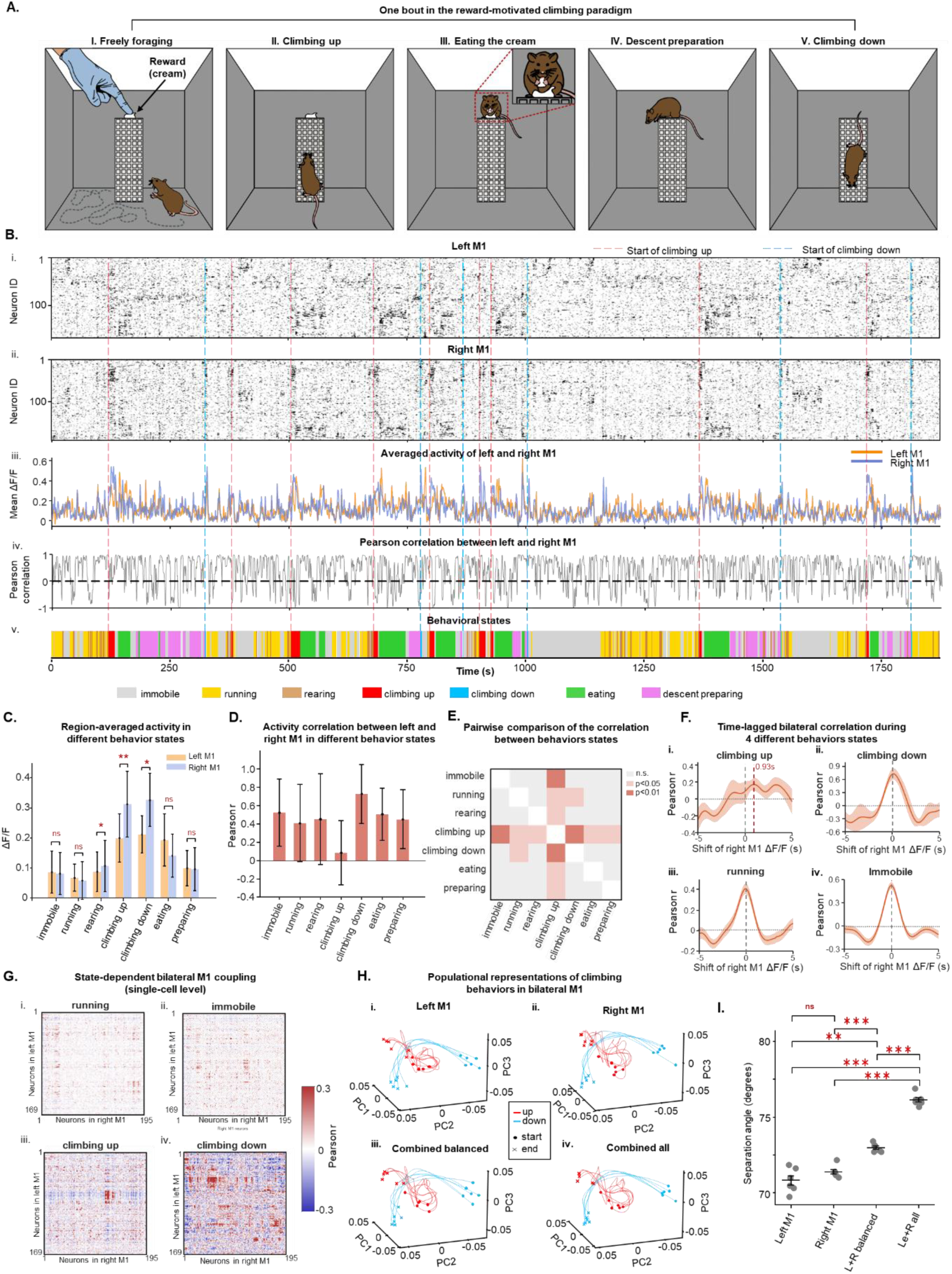
Dual MINI2P-II recordings reveal behavior-dependent inter-hemisphere coordination in bilateral M1 in reward-motivated ladder-climbing tasks. A) Cartoon illustrating the five phases of one climbing bout in the reward-motivated climbing paradigm. (i) Freely foraging phase. Note that a reward (cookie cream) is placed by the experimenter during this phase. (ii) Climbing-up phase; (iii) Eating phase. Note that the mouse consistently faces the side camera. (iv) Descent-preparation phase; (v) Descent (climbing-down) phase. Note that occasionally the mouse will jump from the platform rather than descend via the ladder. B) Behavior-dependent population activity in left and right M1. (i) Left M1 raster (169 neurons) sorted with Rastermap to visualize shared activity motifs; pairwise similarity decays smoothly with pairwise distance in the sorting. Color encodes ΔF/F (darker = higher activity). (ii) Right M1 raster (all 195 neurons), same sorting procedure. (iii) Session-wide mean ΔF/F for left (orange) and right (blue) M1 over ∼29 min. (iv) Sliding-window Pearson correlation (4 s window, 1-frame step) between the left and right M1 population mean ΔF/F traces shown in B-iii. Values range from −1 (anti-correlated) to +1 (co-activated). (v) Manually annotated behavioral epochs: run (yellow), rearing (brown), climb-up (red), climb-/jump-down (blue), eating (green), pre-descent preparation (pink), and inactive/rest (including grooming, quiet wakefulness, sleep, gray). Red dashed lines indicate the starting time point of 9 climbing trials (6 successful, and 3 aborted before reaching the top), while blue lines indicate the starting time point of climbing down. C) State-dependent activity averages. Mean ΔF/F across all neurons in left (orange) and right (blue) M1, grouped by behavioral state. Paired t-test, n.s., non-significant; *, p < 0.05; **, p < 0.01. D) Distribution of sliding-window bilateral correlation values (4 s window) grouped by behavioral state. Bar height, mean correlation; error bar, SD. Note the significantly lower bilateral correlation during climbing-up compared to other states. E) Pairwise statistical comparison of bilateral correlation across behavioral states (Wilcoxon rank-sum test). Core in each cell encodes significance level: light gray, not significant (p ≥ 0.05); faded pink, p < 0.05; medium red, p < 0.01; dark red, p < 0.001. Diagonal cells are white. F) Time-lagged cross-correlation analysis. The mean population ΔF/F of right M1 was shifted relative to left M1 in steps of one frame (1/7.5 s) over a range of ±5 s, and the sliding-window Pearson correlation (4 s window) was recomputed at each lag. Line, mean correlation; shaded area, ±SEM. Dashed vertical gray line, zero lag; dashed horizontal line, zero correlation. (i) Climbing-up. Note the peak at ∼0.93 s lag, indicating that right M1 activity leads left M1. (ii) Climbing-down. (iii) Running. (iv) Immobile. In ii–iv, the peak is centered near zero lag, indicating temporally aligned bilateral activity. G) Neuron-by-neuron cross-hemispheric Pearson correlation matrices between all left M1 and right M1 neuron pairs, computed within each behavioral state. Neurons are reordered by hierarchical clustering (Ward’s method) applied to the climbing-up correlation matrix; the same ordering is used across all panels to enable direct comparison. Color scale, Pearson r (blue, negative; white, zero; red, positive; range [−0.3, 0.3]). (i) Running. (ii) Immobile. (iii) Climbing-up. (iv) Climbing-down. H) 3D state-space trajectories for Left (i), Right (ii), combined balanced (iii), and combined all (iv) neural ensembles. Similar neuron counts were used for i-iii (n=150) but larger in iv (364, 169 Left + 195 Right). Red and cyan lines represent individual climbing up and climbing down trajectories. Start and end points are indicated by dots and crosses, respectively. I) Scatter plot with mean ± SD bars representing the separation angle (in degrees) between individual climbing up & down trajectories in 15-D state space. Paired t-test, ns, non-significant; *, p < 0.05; **, p < 0.01; ***, p < 0.001.

In each experiment, the mouse was first placed on the floor of the arena to forage freely (foraging phase, **Fig. 3A-i**). Three behavioral states were manually annotated by reviewing both side and top camera recordings: immobility (grooming, quiet wakefulness, or sleep; **Fig. S3A-i**), running (**Fig. S3A-ii**), and rearing (**Fig. S3A-iii**). During this phase, the experimenter placed a small piece of cookie cream on the front edge of the top of the platform as a reward (**Fig. 3A-iv**). The mouse then climbed up the ladder (climbing-up phase, **Fig. 3A-ii** and **Fig. S3A-v**). Upon reaching the top, the reward position at the platform edge required the mouse to turn its head toward the side camera while eating it (eating phase, **Fig. 3A-iii** and **Fig. S3A-vi**). After eating, the mouse prepared to descend by placing its forelimbs over the platform edge and repeatedly lowering and retracting its head (descent-preparation phase, **Fig. 3A-iv** and **Fig. S3A-vii**), before descending the ladder (climbing down, or so-called descent phase, **Fig. 3A-v** and **Fig. S3A-viii**). Occasionally, the mouse would jump from the platform rather than climb down. These five phases together defined one complete climbing bout. Once the animal returned to the floor, a new bout began with a fresh reward placed on the platform immediately. This paradigm thus captures a rich diversity of behavioral states, spanning immobile states (grooming and quiet wakefulness), stereotyped locomotion (running and rearing), and bilaterally coordinated actions (eating, climbing up and descending).

To control for regional variability in transgene expression and activity levels, we first ensured that the recording fields were in the forelimb motor area in both hemispheres (*27*). We did this in each mouse by injecting a retrogradely expressing virus (rAAV2-CAG-tdTomato) in a single region in left M1 (coordinates: AP +1.0 mm, ML −1.8 mm, DV 350 µm below the surface) corresponding to the forelimb subregion (*27*). Histological analysis showed a clear and spatially restricted topographical projection between left and right M1, consistent with established anatomical studies(*27*) (**Fig. S3B**). We ensured that all 2P imaging FOVs were positioned within the area of the contralaterally projecting tdTomato-labeled M1 neurons visible in the red channel of each MINI2P-II miniscope (**Fig. S3C**).

#### ii. Behavior-dependent bilateral M1 engagement

In one example ladder-climbing experiment (**Fig. 3B**), we first clustered population activity with Rastermap (*28*), allowing visualization of where and when patterned ensemble activity occurred in each hemisphere (**Fig. 3B-i** left M1, 169 neurons; **3B-ii** right M1, 195 neurons; neurons were from the same recording as in **Figs. 2K** and **2L** with removing neurons with low SNR and few number of calcium transients). We then computed the mean ΔF/F in each hemisphere (**Fig. 3B-iii**), as well as a sliding-window (5 s) Pearson correlation between the two mean traces (**Fig. 3B-iv**) and aligned these signals to frame-by-frame behavioral annotations (**Fig. 3B-v**). Both hemispheres showed significantly different mean ΔF/F across the seven behavioral states (**Fig. 3C** and **Fig. S3D**; Kruskal–Wallis test), with increased activity during climbing-up, climbing-down, and eating, and decreased activity during running (pairwise comparisons in **Fig. S3D**). This pattern of reduced bilateral activity during stereotyped locomotion but increased activity during movements demanding coordination is consistent with previous findings that inter-hemispheric communication is actively recruited when bilateral limb coordination is required (*29*). We also compared hemispheric differences within each state (**Fig. 3C**) and found that right M1 was significantly more active than left M1 during climbing-up, climbing-down, and rearing, but not during the remaining states (Wilcoxon signed-rank test, climbing-up, p= 0.0039; climbing-down, p= 0.0312; rearing, p= 0.0190). This indicates that bilateral recordings can capture behavior-dependent lateralization that would be invisible from one side alone.

#### iii. Behavior-dependent cross-hemispheric correlation at the population level

Next, we investigated whether bilateral M1 coupling varies across behavioral states by computing the mean Pearson correlation between the left and right population mean ΔF/F within each state (**Fig. 3D**). In most states, bilateral correlations were positive and of similar magnitude (**Fig. 3D** and **3E**), indicating a general tendency toward co-activation across hemispheres. However, bilateral correlation during climbing-up was significantly lower than during other states including climbing-down and eating, even though all three require bilateral coordination (**Fig. 3D**; pairwise comparisons, **Fig. 3E**, p-values of the comparisons). To investigate whether this reduced coupling reflected a general decrease in co-activation or the presence of temporally offset rhythmicity between hemispheres, we performed a time-lagged cross-correlation analysis. Specifically, we shifted the mean population ΔF/F of right M1 relative to left M1 in steps of one frame (1/7.5 s) over a range of ±5 s and recomputed the sliding-window Pearson correlation (4 s window) at each lag. For each climbing-up bout, the mean correlation was extracted at every lag, yielding a lag-correlation profile averaged across bouts (**Fig. 3F**). The correlation peaked at a lag of approximately 0.93 s (**Fig. 3F-i**), indicating that right M1 activity consistently preceded left M1 during climbing-up. In contrast, other states, e.g. climbing-down, running, and immobility (**Fig. 3F-ii** to **iv**) all peaked near a zero lag, indicating temporally aligned bilateral activity. This temporal asymmetry during climbing-up may suggest a sequential bilateral organization across left and right M1 during a unique whole-body movement that would not be evident in unilateral recordings.

#### iv. Cross-hemispheric correlation at single-cell resolution

The population-level analyses above treated each hemisphere as a single entity by averaging across all neurons. However, bilateral coordination during complex behaviors may involve specific neuronal subsets rather than uniform hemispheric co-activation — a level of detail that requires simultaneous single-cell resolution in both hemispheres. To investigate this, we computed neuron-by-neuron Pearson correlation matrices between all left and right M1 neuron pairs within each behavioral state and applied hierarchical clustering (Ward’s method) to reveal cross-hemispheric cell assembly structure (**Fig. 3G** and **Fig. S3E**). During most states (**Fig. 3G-i**, running; **Fig. 3G-ii**, immobile; **Fig. S3E**, rearing, eating, and descent preparation), the correlation matrices were relatively uniform, with broadly distributed weak positive correlations. In striking contrast, the climbing states exhibited far more complex structures (**Fig. 3G-iii** and **iv**). During climbing-up, a distinct cluster of neurons in left and right M1 showed strong positive coupling with each other while being anti-correlated with the remaining population, suggesting that a specific bilateral cell assembly is engaged during this behavior (**Fig. 3G-iii**). During climbing-down, multiple positively correlated clusters coexisted alongside negatively correlated groups (**Fig. 3G-iv**). This rich cross-hemispheric structure was entirely absent in the population-averaged signals, where the mean correlation during climbing simply appeared low. This demonstrates a key advantage of 2P over widefield 1P imaging for bilateral studies: only cellular-resolution recording can reveal the coexistence of positively and negatively coupled subpopulations that underlie the decorrelation apparent at the population-level (*30*).

#### v. Complementary behavioral representations across hemispheres

The analyses above show that bilateral M1 is engaged during climbing with behavior-dependent cross-hemispheric coupling, but with a structure that differs between climbing-up and climbing-down. This raises the question of whether the two hemispheres encode the same or partially complementary behavioral information. To address this, we used dimensionality reduction to visualize neural population trajectories for each hemisphere separately and for the combined bilateral dataset (L+R all) in two climbing states. To avoid sampling bias, the same number of neurons (n = 150) was used when constructing neural state vectors. In addition, we generated a balanced bilateral subsample (L+R balanced) by selecting equal numbers of neurons (n = 75, repeated 100 times with random selection) from each hemisphere, matching the total count of the unilateral datasets. We then fit a principal component analysis (PCA) model to the data and projected each climbing bout into this PCA space (Methods and **Fig. 3H**; only first 3 PCs plotted). Each trajectory represents one climbing bout (six trials total); color indicates behavioral condition (climb-up or climb-down) and start and end points are marked by dots and crosses.

In both individual hemispheres (**Figs. 3H-i** and **3H-ii**) and the combined dataset (**Figs. 3H-iii** and **3H-iv**), trajectories were consistent across trials of the same condition, with start and end points close in state space, while climb-up and climb-down trajectories were clearly separable, indicating that behavioral state (upward vs downward climbing) are encoded by orthogonal neuronal subsets. To quantify this, we compared the mean subspace angles between climbing-up and climbing-down trajectories in 10-dimensional PCA space (**Fig. 3I**). The full bilateral population (L+R all) showed the largest subspace angles (mean ± SD: 76.15± 0.37°; paired t-test, vs left alone p = 4.31 × 10^-5^; vs right alone, p = 4.10 × 10⁻^5^; vs L+R balanced, p = 4.10 × 10^-5^), indicating the greatest geometric separation between behavioral states. Left-only and right-only populations were statistically indistinguishable (70.82 ± 0.74° vs 71.38± 0.36°; p = 0.15), suggesting similar representational capacity. The balanced bilateral subsample showed significantly greater separation (72.98± 0.26°) than either unilateral population (vs left, p = 2.85× 10^-3^; vs right, p = 1.94× 10^-4^), although still lower than the full bilateral dataset, indicating that the left and right hemispheres might together contain information about climbing behavior that is not fully captured when sampling from each hemisphere alone. This increased trajectory separation is not explained solely by increased neuron numbers, as the balanced subsample controls sampling size.

Together, these results illustrate that simultaneous bilateral M1 recordings with cellular resolution can reveal behavior-dependent cross-hemispheric functional coupling at the single-cell level, as well as partially complementary representations at the population level. These features are inaccessible to unilateral recordings, lower-yield methods such as electrodes, or population-level bilateral methods such as widefield imaging or fiber photometry. Further work will be needed to fully characterize the functional significance of these inter-hemispheric dynamics and their roles in specific movement phases (e.g., initiation, stable climbing, and transition), which is beyond the scope of the present study (see Discussion).

### 2P imaging in two ipsilateral brain regions simultaneously in freely moving mice

#### i. A new implantable optical assembly enables imaging of adjacent cortices with two MINI2P-II miniscopes

So far, we have shown that MINI2P-II supports dual-region imaging across widely separated brain areas. However, a limitation here is that the two regions must be at least a few mm apart to prevent physical collision of the two miniscopes, preventing many applications that require imaging of anatomically adjacent brain structures.

To address this, we developed a novel “periscope” optical assembly termed the GRIN periscope assembly (GPA), inspired by periscope lenses in recent smartphones (*31*), that shifts the imaging FOV both vertically and laterally away from the recording region while preserving imaging resolution and magnification (**Fig. 4A**). Each assembly consists of two 1-mm diameter GRIN lenses, two microprisms and a coverslip, with a step-by-step protocol provided in **Fig. S4A**. When two GPAs are implanted side by side, the FOV centers are as close as 1 mm in the tissue but the miniscopes are 8.4 mm apart, compatible with the 5-mm footprint of the MINI2P-II (**Fig. 4B**). This geometry allows two miniscopes to be mounted simultaneously without collision when imaging nearby regions. The vertical offset also permits imaging of deep targets such as hippocampal CA1.

**Figure 4.**
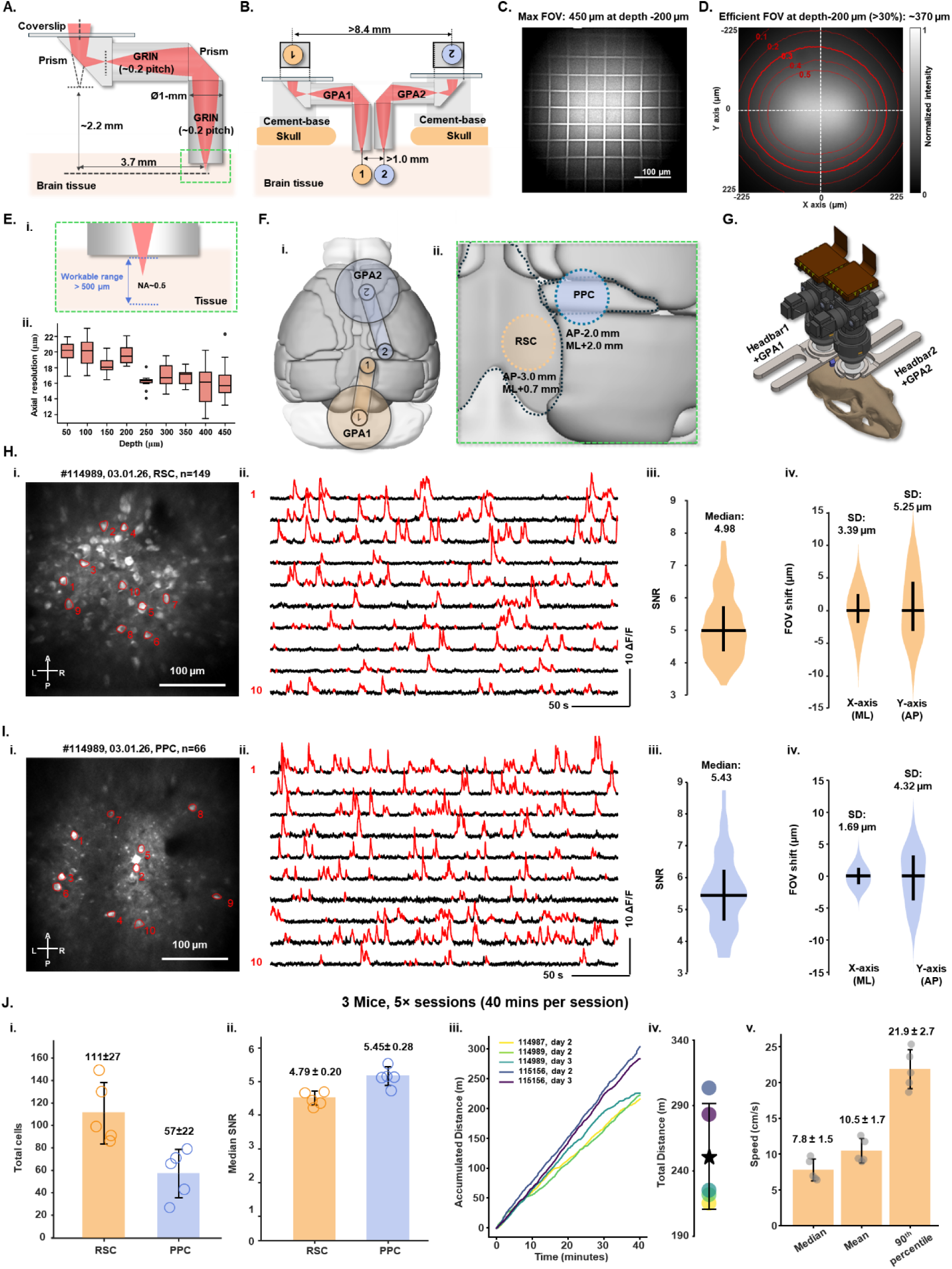
GRIN periscope assembly (GPA) enables dual-miniscope imaging of adjacent brain regions. A) Optical layout of a single GRIN periscope assembly (GPA). The GPA consists of a 0.15-mm-thick coverslip, a 1.3-mm-wide prism, a 0.2-pitch GRIN lens, a second 1-mm-wide prism, and a second 0.2-pitch GRIN lens. The excitation light path from the objective to the sample is shown in red, illustrating the periscope-like light bend. B) Two GPAs implanted side-by-side for adjacent-region imaging. Although the actual cortical targets are ∼1 mm apart in tissue, the GPA shifts the two FOV centers to >8.4 mm apart, exceeding the 5-mm MINI2P-II footprint and preventing collision. The circles 1 and 2 indicate the two FOVs, with mirror inversion introduced by the GPAs. Cement bases are fixed to the skull to stabilize both GPAs. C) Grid imaging through one GPA. MINI2P-II imaging of a 50-µm grid at 200 µm depth demonstrates a maximum FOV of 450 × 450 µm, confirming 1:1 magnification. Note the reduced intensity at the edges, indicating the efficient FOV is smaller than the maximum FOV (region supporting high-SNR calcium signals). D) Homogeneity and effective FOV. Imaging a uniform fluorescent slide at 200 µm depth shows an effective FOV about 370 µm (diameter), defined as the region retaining ≥30% of peak intensity. Red contours indicate 50%, 40%, 30%, 20%, and 10% levels. E) Axial resolution maintained across depth. (i) Illustration of the designed numerical aperture (NA) of the GPA, and the 0–500 µm depth range for axial resolution measurement ((ii) and **Fig. S4B**). (ii) Axial resolution measured using 1-µm fluorescence beads embedded in 1.5% agarose solution in the ratio of 1:100 in volume. F) Cortical targeting of two GPAs (dorsal view). (i) GPA1 targets retrosplenial cortex (RSC) (orange dashed circle) and GPA2 targets posterior parietal cortex (PPC) (blue dashed circle) in right hemisphere. (ii) Zoom-in with coordinates: RSC (AP −3.0 mm, ML +0.7 mm); PPC (AP −2.0 mm, ML +2.0 mm). G) Mounting two miniscopes onto GPA implants. Each miniscope is aligned to its own headbar. H) Example imaging from RSC. (i) Maximum-intensity projection from the RSC FOV; 149 neurons detected. (ii) Ten example ΔF/F traces from neurons marked in h-I (red parts show suprathreshold transients). (iii–iv) Violin plots of signal-to-noise ratio (SNR) for all detected neurons (n =149) and rigid motion (inter-frame FOV drift) along AP (X) and ML (Y) axes. Violin outlines show kernel-density estimates (bandwidth = 10% of data range); vertical lines, interquartile range (IQR); horizontal bars, median. Median SNR and motion SD are indicated on plots (symbols as in I). I) Example imaging from PPC (same session). (i) Maximum-intensity projection from PPC; 66 neurons detected. (ii) Ten example ΔF/F traces from marked neurons in I. (iii–iv). (iii–iv) Violin plots of SNR (n =66) and rigid motion (AP/ML). Plot conventions match h. J) Session-level imaging quality and behavior metrics across animals. Summary for 3 mice, 5 sessions, 40 min each. (i) Total neurons per recording (Mean ± SD): RSC: 111 ± 27, PPC: 57 ± 22. (ii) Median SNR across all recorded neurons for each recording (Mean ± SD, RSC: 4.79± 0.20, PPC: 5.45± 0.28). (iii) Accumulated running distance across each session over 40 min; each line indicates one recording (colored by mouse ID and day). Same color scheme applies to iv. (iv) Total distance traveled across all 5 sessions (mean ± SD shown). (v) Speed metrics across sessions: median, mean, and 90th percentile speed. Bar height, mean; error bar, SD; gray dots, individual sessions; values shown.

The effective FOV, defined as the region that retains at least 30% (∼1/e) of maximum signal amplitude is 370 μm in diameter at the imaging depth of -200 μm (used with MINI2P-II) (**Figs. 4C** and **4D**). The axial resolution at the center of the FOV was stable across a depth range of 50–450 μm (e.g. 19.91 ± 1.57 µm at depth of 50 µm, and 16.37 ± 2.50 µm at depth of 450 µm) (**Figs. 4E, S4B**).

#### ii. Simultaneous 2P imaging in ipsilateral retrosplenial cortex (RSC) and posterior parietal cortex (PPC)

The RSC and PPC are densely interconnected navigation hubs containing large populations of neurons that map local space egocentrically (*32–34*), but how their representations diverge within-animal and during the same behavioral trials is not known. This largely reflects technical limitations: despite their proximity, the two regions cannot be recorded ipsilaterally during real-world 2D navigation with a single miniscope or without multiple spaced recording electrodes or tetrode bundles (*5*).

We demonstrate here how combining GPAs with MINI2P-II miniscopes overcomes this limitation. In three male transgenic mice (CaMKII-tTA/tetO-GCaMP6s), we implanted one GPA over the dorsal surface of the dysgranular RSC in the right hemisphere (center: AP −3.0 mm, ML −0.7 mm) and a second GPA over the medial PPC (center: AP −2.0 mm, ML −2.0 mm) in the same hemisphere (**Figs. 4F**, **4G**, and **S4C**). A MINI2P-II miniscope was then baseplated above each GPA (**Figs. 4G**, and **S4D**).

In an example 40-min recording, a mouse foraged for scattered rewards in an 80-cm arena with a clear visual cue (i.e. a A4 white paper sheet attached on the south wall). We detected 149 highly active neurons in RSC (**Fig. 4H**) and 66 in PPC (**Fig. 4I**), both from a single plane in layer II. Both datasets showed high signal-to-noise ratios (median SNR ± SD: 4.98 ± 1.05 in RSC, **Fig. 4H-iii**; 5.43 ± 1.17 in PPC, **Fig. 4I-iii**) and moderate motion artifacts (Figs. **4H-iv**, **4I-iv**; **Video S3**). The lower yield in PPC thus more likely reflects variation in GCaMP expression rather than imaging performance.

Across three mice and five 40-min sessions, we recorded a total of 555 neurons from RSC and 286 from PPC. RSC consistently yielded more cells (**Fig. 4J-i**), whereas SNR was similar between regions in all mice (**Fig. 4J-ii**). Running distance (**Figs. 4J-iii**, **4J-iv**) and speed (**Fig. 4J-v**) remained stable across all recordings and were comparable to earlier experiments (**Figs. 2E and 2F**). These results indicate that two GPAs combined with two MINI2P-II miniscopes enable high-quality imaging during vigorous behavior and in adjacent cortical regions that were previously inaccessible together.

#### iii. Analyzing egocentric boundary representation in PPC and RSC

Egocentric boundary cells (EBCs), which are abundant in both RSC and PPC, fire when environmental boundaries appear at a specific orientation and distance relative to the animal (*35–37*). Using our dual-region recordings in freely moving mice, we asked whether EBCs in the PPC or RSC exhibit similar or distinct functional properties within the same animals and during the same behavioral sessions.

To identify EBCs, we constructed egocentric boundary rate maps (EBRs) following established procedures (*35*) (**Fig. S5A**; Methods). Briefly, for each video frame, we computed the distance to the nearest wall at every 3° offset relative to the animal’s head direction. We applied the same procedure to each deconvolved spike event for all neurons. EBRs were then generated in polar coordinates by dividing spike counts by occupancy time in each angular–distance bin. From each EBR, we computed the mean resultant length (MRL) of angular tuning for the full session. We also quantified the absolute differences in preferred direction and preferred distance between the first and second halves of the recording. A neuron was classified as an EBC if it satisfied three criteria derived from prior work (*35*): (i) its session MRL exceeded the 99th percentile of a shuffled distribution created by circularly shifting its spike train; (ii) its preferred direction changed by <45° between session halves; and (iii) its preferred distance differed by <50% from the full-session estimate in both halves.

Neurons that met these criteria showed sharp egocentric boundary tuning in both regions. **Figs. 5A** to **5C** illustrate three examples from the RSC, and **Figs. 5E** to **5G** show three examples from the PPC from same recording. As in previous studies, we observed three types of EBCs in both areas. The first type fired when the animal was very close to boundaries at a specific orientation (animal-proximal fields; **Figs. 5A** and **5E**). The second type fired at farther distances from boundaries (animal-distal fields; **Figs. 5B** and **5F**). The third type decreased activity when the animal was close to the boundaries (inverse EBCs; **Figs. 5C** and **5G**). The fractions of different types of EBCs for the example session are shown in **Fig. 5D-i** (RSC) and **Fig.5H-i** (PPC). These results demonstrate that our dual-scope recordings can resolve fine-scale egocentric tuning features in both cortical areas (Figs. 5D-i).

**Figure 5.**
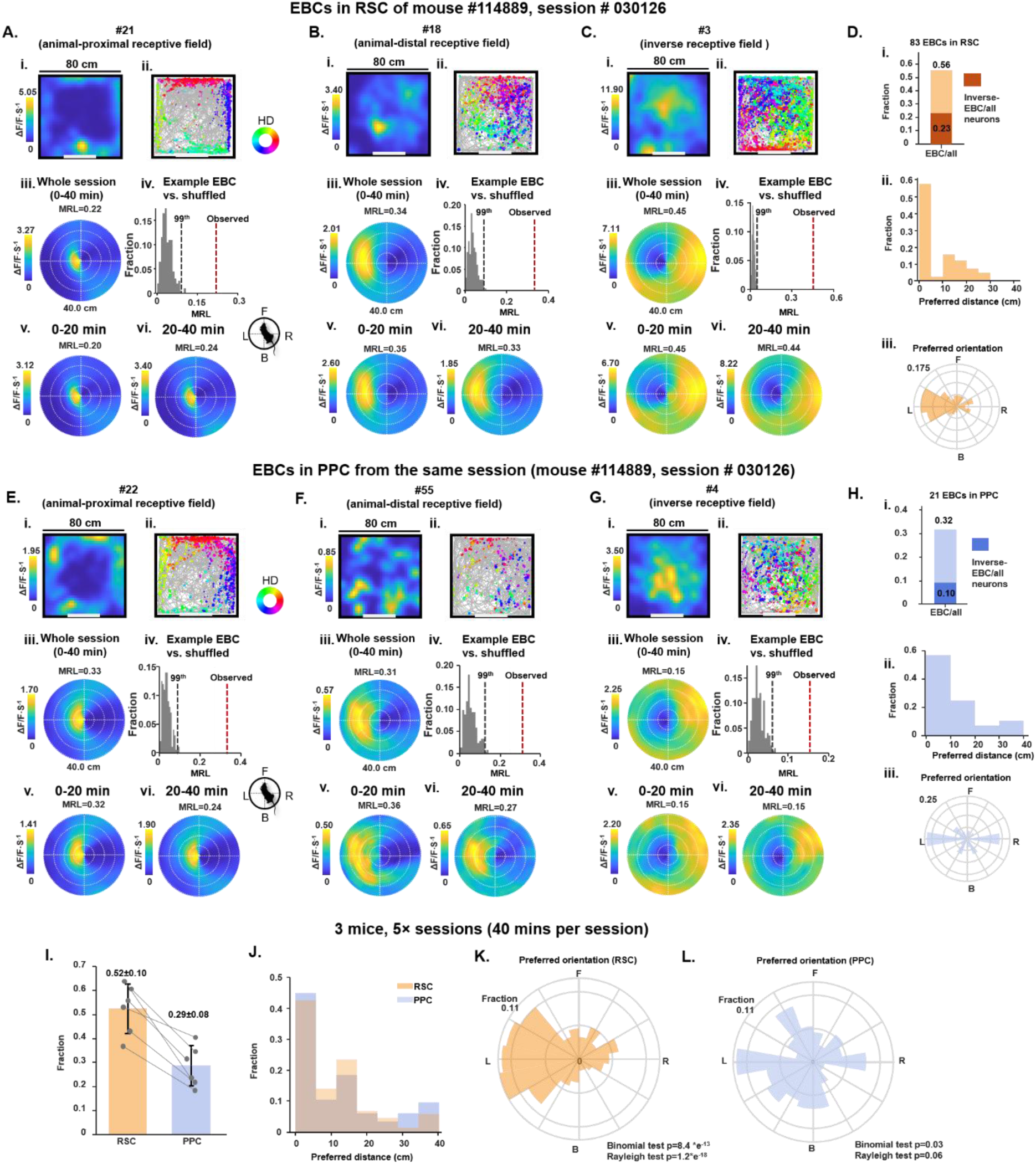
Functional properties of egocentric boundary cells (EBCs) in PPC and RSC. A) RSC EBC example 1—animal-proximal receptive field. (i) spatial tuning of the example EBC in the full 40 min session, displayed as intensity of calcium activity (ΔF/F per second, color scale). Spatial bins: 2.5 × 2.5 cm^2^; 3 cm Gaussian smoothing kernel. Top numbers indicate peak activity (ΔF/F·s⁻¹). (ii) mouse trajectory (in gray) with calcium events superimposed (same cell as in i). Events are color-coded by head direction; size of spots indicates amplitude. (iii) egocentric boundary rate map (EBR) for the full session showing calcium activity with respect to egocentric boundary locations in polar coordinates. The mean resultant length (MRL) is shown. (iv) MRL of the example EBC (red bar) exceeds 99th percentile (blue bar) of shuffled data (gray bars, 200 iterations). (v–vi) EBRs for the first and second 20-min halves. Note the localized high-activity “bump” close to the head on the left side in full and split-session maps. B) RSC EBC example 2—animal-distal receptive field. (i–vi) Conventions as in A. Note the localized bump at a farther egocentric distance on the left side across full and split sessions. C) RSC EBC example 3—inverse receptive field. (i–vi) Conventions as in A. Note a localized negative bump (decreased activity) to the left of the head across full and split sessions. D) RSC statistics for example session (mouse #114889, session #030126). (i) Fractions of all EBCs, and inverse EBCs in the all recorded RSC among all recorded neurons (n = 149); total EBCs = 83. (ii) Distribution of preferred distance for all 83 RSC EBCs. (iii) Polar histogram of preferred orientation across the same 83 EBCs (top-left number indicates the maximum fraction bin). Note the predominance of leftward receptive fields. E) PPC EBC example 1—animal-proximal receptive field (same session as **A-D**). (i–vi) Conventions as in a. F) PPC EBC example 2—animal-distal receptive field. (i–vi) Conventions as in **B**. G) PPC EBC example 3—inverse receptive field. (I–VI) Conventions as in **C**. H) PPC statistics for the same example session. (i) Fractions of All-EBC, Normal-EBC, and inverse EBC among all recorded neurons (n = 66); total EBCs = 21. (ii) Distribution of preferred distance for all 21 PPC EBCs. (iii) Polar histogram of preferred orientation for the same cells (top-left number indicates the maximum fraction bin). I) Cross-session EBC fractions. Fractions of EBCs in RSC and PPC across 3 mice, 5 recordings; gray dots, individual regions per session; gray lines connect the two regions recorded simultaneously. J) Cross-session preferred-distance distributions. Pooled preferred distances for all EBCs in RSC (orange) and PPC (blue) across the 5 sessions (3 mice). K) Cross-session preferred orientations—RSC. Polar histogram showing a strong leftward (contralateral) bias across all RSC EBCs (Binomial p = 8.4 × 10⁻¹³; Rayleigh p = 1.2 × 10⁻¹⁸). L) Cross-session preferred orientations—PPC. Polar histogram showing a more balanced distribution relative to RSC (Binomial p = 0.03; Rayleigh p = 0.06).

In this example dataset (**Figs. 5A** to **5H**), we identified a total of 83 EBCs in the RSC (65%, n = 83/149; **Figs. 5D** and **S5B**) and 21 in the PPC (32%, n = 21/66; **Fig. S5C**). Preferred-distance distributions extended well beyond the range of direct whisker contact in both regions (**Fig. 5D-ii** and **Fig. 5H-ii**). Across all three mice, the fraction of EBCs (i.e., RSC > PPC) and their preferred-distance distributions were consistent (**Figs. 5I, 5J**; **Supplementary Document Part 1**), in agreement with previous reports (*35, 36*).

We next examined the preferred spatial orientations of the cells, in light of all recordings being in the right hemisphere. In the RSC, we observed a strong predominance of leftward (contralateral) receptive fields (**Fig. 5D-iii** for one recording; **Fig. 5K** for all recordings; right-hemisphere EBCs preferring the left spatial hemifield: 68.9%, Binomial p = 8.4 × 10⁻¹³; Rayleigh p = 1.2 × 10⁻¹⁸). In contrast, the distribution of preferred orientations in the right PPC was more balanced between left- and rightward fields (**Fig. 5H-iii** and **Fig. 5L**; right hemisphere EBCs encoding the left side: 60.6%, Binomial p = 0.03; Rayleigh p = 0.06). The distributions of preferred orientations differed significantly between RSC and PPC EBCs (Watson’s two-sample U² test, U² = 0.205, p = 0.0346, permutation test with 100,000 iterations). These differences suggest a previously unrecognized regional difference that we were able to resolve with MINI2P-II and the GPA assemble, in which the RSC more strongly represents contralateral egocentric boundaries, whereas the PPC encodes egocentric space more evenly and may contain proportionally fewer EBCs, at least in superficial cortical layers.

##### Populational 2P imaging in freely moving adult and juvenile zebra finches

The ultralight weight of MINI2P-II makes it possible to perform calcium imaging in smaller species than mice and at earlier developmental stages than previously accessible with 2P miniscopes. Zebra finches, weighing only 13–16 g in adulthood, are a well-established model for studying motor learning (*38*), plasticity and sequence coding (*39*) (**Fig. 6A**, left). Juvenile male finches (**Fig. 6A**, right) undergo song learning during a well-defined sensitive period that spans roughly 25–90 days post-hatching (dph, **Fig. 6B**) (*40*), during which the birds grow from about 6 grams to their eventual adult weight (*41*). These lightweight animals are highly sensitive to head-mounted loads; excess weight causes stress and disrupts natural song learning and singing behaviors (*42, 43*). Accordingly, existing 2P miniscopes (>2 g) require extensive training or habituation for use during singing, a limitation that is even more pronounced in lightweight juveniles during natural sensory learning stages (<50 dph, **Fig. 6B**).

**Figure 6.**
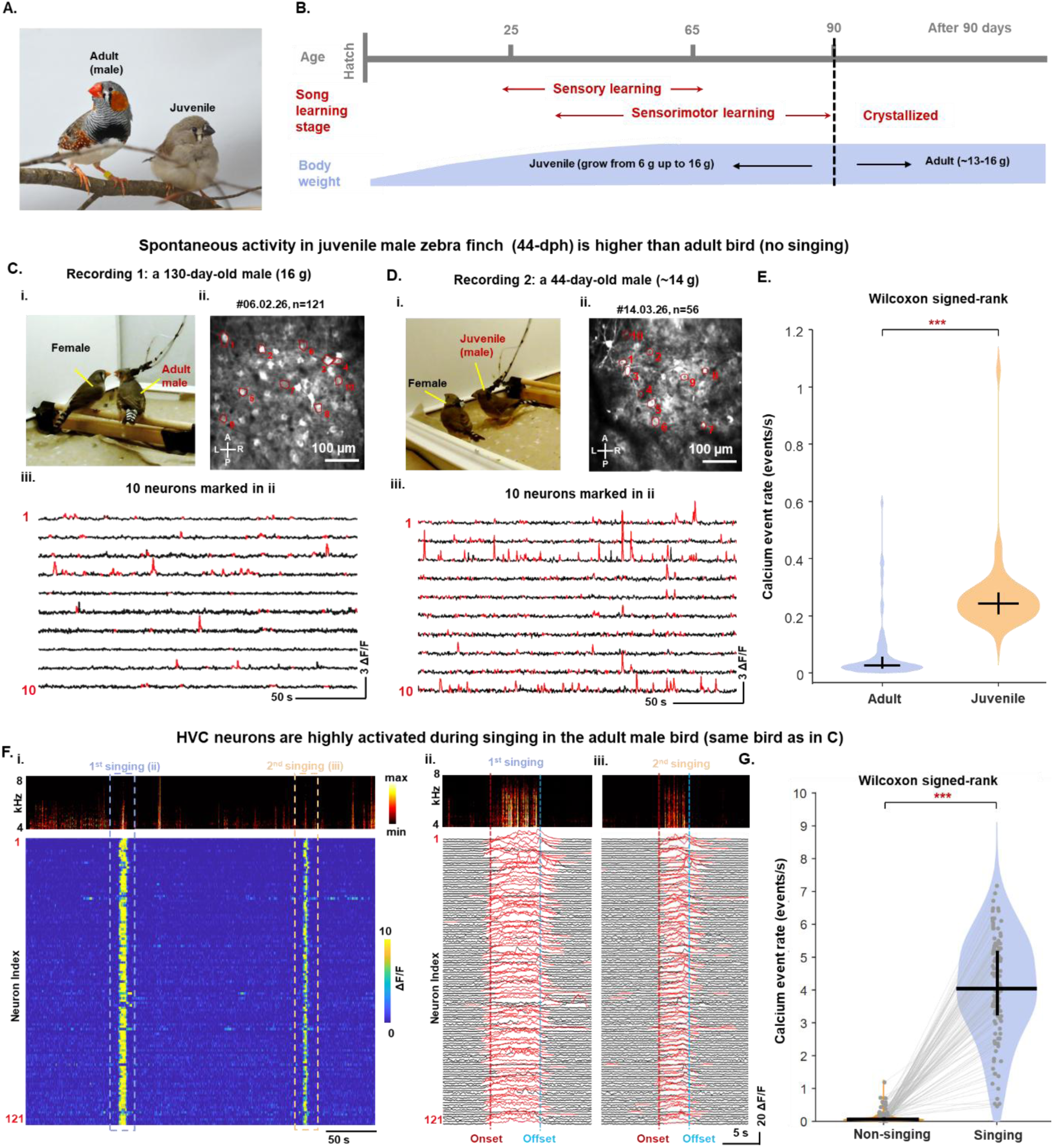
Calcium imaging in adult and juvenile zebra finches. A) Photographs of an adult male zebra finch (left) and a ∼30 dph juvenile (right), shown side-by-side prior to virus injection and surgery. B) Developmental timeline of song learning and body-mass growth, based on Leitao & Gahr, 2024. Birds are considered juvenile until ∼90 dph (dashed line). Two learning phases overlap: sensory learning (∼25–65 dph; memorization of the tutor song) and sensorimotor learning (onset ∼25–30 dph with subsong babbling; vocal practice guided by auditory feedback). Song typically crystallizes after ∼90 dph (stable adult song). Body weight increases rapidly from hatching to ∼50 dph. C) MINI2P-II recording from a 130-dph adult male zebra finch. (i) Video snapshot of a social interaction session: adult male (right, with MINI2P-II mounted) oriented toward the female (left). (ii) Maximum-intensity projection of the HVC imaging FOV; 121 neurons detected. (iii) Ten example ΔF/F traces from the neurons indicated in c-ii. Red segments indicate suprathreshold calcium transients. D) MINI2P-II recording from a 44-dph juvenile male zebra finch. (i) Video snapshot of a social interaction session: juvenile male (right, with MINI2P-II mounted) oriented toward the female (left). (ii) Maximum-intensity projection of the HVC imaging FOV; 56 neurons detected. (iii) Ten example ΔF/F traces from neurons indicated in **d-ii**. Red segments indicate suprathreshold calcium transients. Note the higher frequency of detectable calcium transients compared with the adult neurons shown in **C-iii**. E) Violin plot comparing spontaneous calcium event rates across all neurons detected in the adult (left) and juvenile (right) birds. Violin outlines show kernel-density estimates; vertical lines indicate the interquartile range (IQR). Event rates are significantly higher in the juvenile than in the adult (Wilcoxon signed-rank: p = 1.44 × 10^-20^). F) HVC activity in the adult male zebra finch is highly song-locked. (i) Top: spectrogram of the sound recording during imaging; two singing periods are indicated (1^st^ singing, 2^nd^ singing). Middle: ΔF/F raster plot across all 121 neurons, sorted by peak amplitude (colormap: yellow, max ΔF/F; blue, min ΔF/F). Note that high activity levels are restricted singing periods, and that activity remains low outside of them. (ii–iii) Zoomed-in views of the 1st and 2nd singing periods from (i), showing the spectrogram (top) and ΔF/F traces of all 121 neurons (bottom). Red segments indicate suprathreshold transients. Onset and offset of each singing period are marked by red and blue dashed lines, respectively. G) Violin plot comparing calcium event rates across all neurons in the adult bird during non-singing (left, yellow) and singing (right, blue) periods. Violin outlines show kernel-density estimates; vertical lines indicate the IQR. Each dot represents one neuron; grey lines connect the same neuron across conditions. Event rates during singing are significantly higher than during non-singing (Wilcoxon signed-rank: p = 1.35 × 10^-21^).

We demonstrate here that MINI2P-II enables large-scale calcium imaging in both adult and juvenile zebra finches. Recordings were obtained from two animals: one adult male (130 days old, 16 g; **Fig. S6A**) and one juvenile male (44 dph, ∼14 g; **Fig. S6B**). Both birds received AAV-GCaMP6s injections (**Fig. S6C**), underwent a craniotomy (**Fig. S6D**), and were baseplated with a MINI2P-II (**Fig.S6E**) targeting the high vocal center (HVC), a premotor area essential for song control. HVC was distinguished from surrounding regions using retrograde labeling of its projection to basal ganglia Area X (**Figs. S6F** and **S6G**) (*44, 45*).

For each experiment, the implanted bird was placed with a female in the same enclosure, allowing them to interact freely (adult: **Fig. 6C**; juvenile: **Fig. 6D**). Both the adult and juvenile males exhibited natural head and body movements and spontaneous social interactions with the female, indicating that the 1.1 g weight of MINI2P-II did not impair motor function regardless of developmental stage (**Video S4**, adult; **Video S5**, juvenile). In the example recordings, we detected 121 neurons in the adult (**Fig. 6C-ii**) and 56 neurons in the juvenile (**Fig. 6D-ii**) within a single FOV, with high spatial resolution and SNR, proving that MINI2P-II could access ensemble activity in zebra finches, and during developmental stages not previously accessible with 2P imaging.

During non-singing behaviors—such as jumping, calling, or interacting with the female—HVC neurons in the adult maintained low spontaneous activity (**Fig. 6C**; **Video S4**). By contrast, HVC activity in the juvenile bird was substantially higher (**Fig. 6D**; **Video S5**). Quantification of calcium event rates across all recorded neurons confirmed this difference (**Fig. 6E**; Wilcoxon rank-sum: p = 1.44 × 10^−20^).

In the adult experiment, within only a few interactions the adult male initiated natural courtship behavior—circling and jumping around the female, nodding his head, gently touching her with his beak, and ultimately singing while oriented toward her (**Video S4**), reflecting strong motivational drive for directed singing. Since HVC activity in adult males is highly song-locked (*42*), we expected to see activity increase dynamically while the adult bird sang. Indeed, during two singing periods in the experiment (**Fig. 6F**; singing period 1: **Fig. 6F-i**; singing period 2: **Fig. 6F-ii**), nearly all recorded HVC neurons showed robust increases in activity relative to non-singing periods (**Fig. 6G**; Wilcoxon signed-rank: p = 1.35 × 10^−21^). This contrasted with the higher-level of spontaneous HVC activity in the juvenile bird, potentially reflecting a more dynamic state of the premotor system during development and the active sensorimotor learning phase.

Together, these proof-of-concept experiments demonstrate that MINI2P-II enables stable, high-quality, large-scale calcium imaging in freely interacting adult and juvenile zebra finches. This provides a powerful platform for studying learning, circuit development and social behavior across previously inaccessible developmental stages when natural learning occurs.

## Discussion

A central challenge in systems engineering is to reduce the size and weight of a complex instrument without sacrificing its performance. This challenge is especially important for head-mountable neural-recording tools in neuroscience, which require high throughput (high cell yield, wide FOV and fast acquisition), high resolution (cellular and subcellular), and minimal weight so that model animals, typically rodents and birds, behave naturally during recording. Miniature 2P microscopes are an excellent example of engineering innovation responding to this need (*21*). After 25 years of iteration since their first prototype in 2001 (*12*), their optical performance now approaches benchtop systems while weighing nearly 5,000× less (≈2 g vs. >10 kg). Reducing weight yet further, without losing speed, resolution or FOV, has been difficult because most attempts have targeted individual components while leaving the optical architecture largely unchanged. For example, using flat lenses (e.g. meta-lenses) to replace the traditional multi-element objective reduced the total weight of the 2P miniscope to near 1 gram (*22*). But the strong angle-dependent aberration and low efficiency of meta-lenses decreased the spatial resolution, limited field of view and reduced the detection sensitivity. Other studies simplified the miniscope structure with removing optical elements such as scan lens and focus-tunable lenses (Tlenses) (*23*), or the entire collection optics with de-scanning schema (*46*), again with a trade-off of poor spatial resolution, no z-scanning capability, or low efficiency.

Here, we step beyond the state of the art by demonstrating several new optical design principles. The first is to reuse optical elements for multiple functions whenever their specifications are the same or similar. Second, integrate components which can be optically congregated into a single device. Third, fold the static-excitation, scanning-excitation and emission paths into the same physical volume to maximize spatial compression. By following these principles and developing new optical components, we reduced the miniscope weight by more than half without sacrificing imaging performance. Further weight reduction while maintaining the current image quality might be achieved by incorporating low-aberration flat optics (e.g., high-reflection-index metamaterials (*47*)), a single-element 3D scanner (*48*) and lighter optical components (using thin glass to make the BFM). This miniscope architecture also creates room for integration of more complex functions, such as single-cell stimulation (*23*) with a miniature spatial light modulator (*49*) or a higher-NA (typically heavier) objective for improved spatial resolution, while remaining under the behavior-compatible weight limit of 3 g for mice. Equally important, the modular design of MINI2P-II should accelerate technological iterations by allowing module-specific upgrades or additions without re-designing the entire miniscope.

Beyond delivering a lighter device, this study provides comprehensive, ready-to-use solutions for multi-region imaging with multiple new miniscopes including a new implantable optical element—the GRIN Periscope Assembly (GPA). Beyond allowing imaging of adjacent regions, the GPA also enables access to regions where mounting a miniscope is typically difficult, such as extreme anterior areas (e.g., olfactory bulb), lateral areas (e.g., auditory cortex, lateral entorhinal cortex), and posterior sites (e.g., cerebellum), by shifting the imaging field to positions where the miniscope can be mounted more easily (e.g., near the skull center). Even within head-fixed paradigms, multiple GPAs with multiple MINI2P-IIs can be implanted to access a larger number of brain regions simultaneously, without the restriction of mounting weight. A current limitation of the GPA is the restricted effective FOV (and associated cell yield) due to optical aberrations introduced by GRIN lenses; this may be mitigated by adding correction elements such as compensation lenses (*50*). Another limitation is that the minimum distance between two regions is 1 millimeter, still larger than the 450-µm FOV that one MINI2P-II can cover. Recently developed large-FOV 2P miniscopes may cover this gap (*20*).

In the bilateral M1 experiment, we used the ladder-climbing task to demonstrate that MINI2P-II can capture inter-hemispheric dynamics at multiple levels of resolution. At the population level, we observed behavior-dependent changes in bilateral coupling strength and temporal alignment, including demonstration of right M1 leading left M1 during climbing-up epochs that was absent during other states. At the single-cell level, hierarchical clustering of neuron-by-neuron cross-hemispheric correlation matrices revealed structured cell assemblies during climbing that were invisible in population averages. At the population-geometry level, combining neurons from both hemispheres improved the separability of climbing-up and climbing-down trajectories beyond what either hemisphere achieved alone. These observations, obtained from a single example session, provide a proof-of-concept demonstration of the analytical possibilities enabled by dual-miniscope bilateral imaging. However, whether cross-hemispheric coordination plays a critical role in controlling such complex full-body movement remains unclear. This question could be investigated by adding an optogenetics module into MINI2P-II (**Fig. S1F**) to perform localized perturbations in one or both hemispheres at different phases of climbing, including initiation, stable climbing and top-reaching, while monitoring effects locally, across hemispheres, and on behavior. Furthermore, by leveraging the capability of MINI2P-II for long-term, single-neuron tracking, it will be possible to examine whether and how such cross-hemispheric interactions evolve during learning or skill acquisition (e.g., improved climbing performance across days).

In a separate experiment, we examined egocentric boundary representations in ipsilateral RSC and PPC. At the population level, we observed similar but not identical egocentric boundary representations in these two regions. It remains unclear whether the deviation reflects differences in inputs or local circuit computations in these two regions. Future studies could take advantage of dual-color MINI2P-II imaging combined with retrograde tracing (such as in **Fig. S3B**) to dissect the functional properties of projection-defined neuronal populations projecting to and between these regions, thereby addressing this question.

Finally, the ∼1 g mass of MINI2P-II makes it possible to image large population of neurons in very small animals and at young ages. The zebra finch is an excellent example of this, since free movement and social interactions are imperative for vocal learning, yet existing tools impose a substantial weight load on these lightweight birds. Wire electrode recordings in freely singing juvenile zebra finches have been achieved as early as 38 dph (*51*), but electrodes typically capture a small number of neurons at a time. High-density electrophysiological recordings, such as with Neuropixels, have not been established in freely moving juvenile finches, because the combined weight of the probe and protective head cap required for chronic use is impractical at these body weights. The only previous calcium imaging in a freely-moving juvenile finch, i.e., a 1P miniscope in a 49-dph bird, provided proof-of-concept results (*43*), but with limited spatial resolution and poor signal-to-noise ratio. MINI2P-II overcomes all of these limitations at once: it records far more neurons than wire electrodes, weighs less than chronic implants for silicon-probes, and produces much higher SNR and spatial resolution than 1P miniscopes. This enabled us to image tens of clearly resolved neurons in a freely moving 44-dph juvenile finch, representing the youngest stage at which population- and cellular-resolution recordings have been reported in a freely behaving songbird. By further optimizing viral expression and surgical protocols, we aim to push imaging to earlier developmental stages with lower body weight, ultimately covering a broader range of the song-learning period. The 2P access to early developmental windows when animals are too small or weak to carry heavier devices also enables reliable longitudinal tracking of the same neurons across development, which is particularly valuable for probing circuit wiring and functional maturation. We anticipate that mouse and rat pup studies will also benefit from this advantage in coming years.

Overall, MINI2P-II, together with practical solutions for multi-region and multi-species imaging, sets a new standard for instrument miniaturization and provides a powerful toolset for investigating neuronal computation during unrestricted behaviors across spatiotemporal scales, developmental stages and species.

## Acknowledgments

We thank L. Bierbusse, M. R. Jorge, A. Foroughi, and P. I. Andersen for technical assistance; E.I. Moser, M.-B. Moser, L. Luo, H. Wu, H. Kanodia, and S. Arber for comments and discussions; W. Tang for viral reagents; P. Dupriez for laser development; P. Kolsch for polarization-maintaining fiber development; and M. S. Pofahl and M. A-Haug for surgical suggestions. This work was supported by the Research Council of Norway FORSKER25 grant (ES758698, W.Z.), FRIPRO grant (353808, W.Z.), Proof of Concept grant (355770, W.Z.), Centre of Excellence grant (Centre for Algorithms in the Cortex 332640, W.Z.), National Infrastructure grant (NORBRAIN IV, grant 350201, W.Z.) and Open access funding provided by the Max Planck Society (M.G).

## Author contributions

Conceptualization: W.Z. Supervision: W.Z., M.G. System design: W.Z. Hardware development: W.Z., K.J., E.K-S. Software development: W.Z., A.L.C. System characterization: E.K-S., W.Z. Mouse surgery and experiments: W.Z. Zebra finch surgery and experiments: S.M., W.Z. Behavior influence tests: W.Z, J.A., E.K-S. Data analysis: E.K-S., S.M., J.A, R.S., W.Z., J.W. Figures: W.Z., E.K-S., S.M., J.A., R.S. Videos: W.Z., E.K-S., S.M. Writing – original draft: W.Z. Writing – review & editing: J.W., E.K-S., S.M., M.G. Discussion and comments on paper draft: all authors.

## Declaration of interest

We are applying for patents under Patent Cooperation Treaty (PCT) Application Nos: [PCT/EP2026/059476 and PCT/EP2026/059496]. The MINI2P-II system is freely available for non-commercial academic use, including duplication and modification. Commercial use is subject to the terms of the applicable patent protection.

## Data and code availability

The data for each figure in this manuscript will be available as of the date of publication. Any additional information required to reanalyze the data reported in this paper is available upon request. 3D models and drawings of the customized components, building instructions and video tutorials, and analysis code, can be found here: https://github.com/kavli-ntnu/MINI2P_toolbox. The original, unprocessed data will be published through Mendeley Data (DOIs: 10.17632/8fjcwfn3mv.1) of the date of publication

---

**Figure S1.**
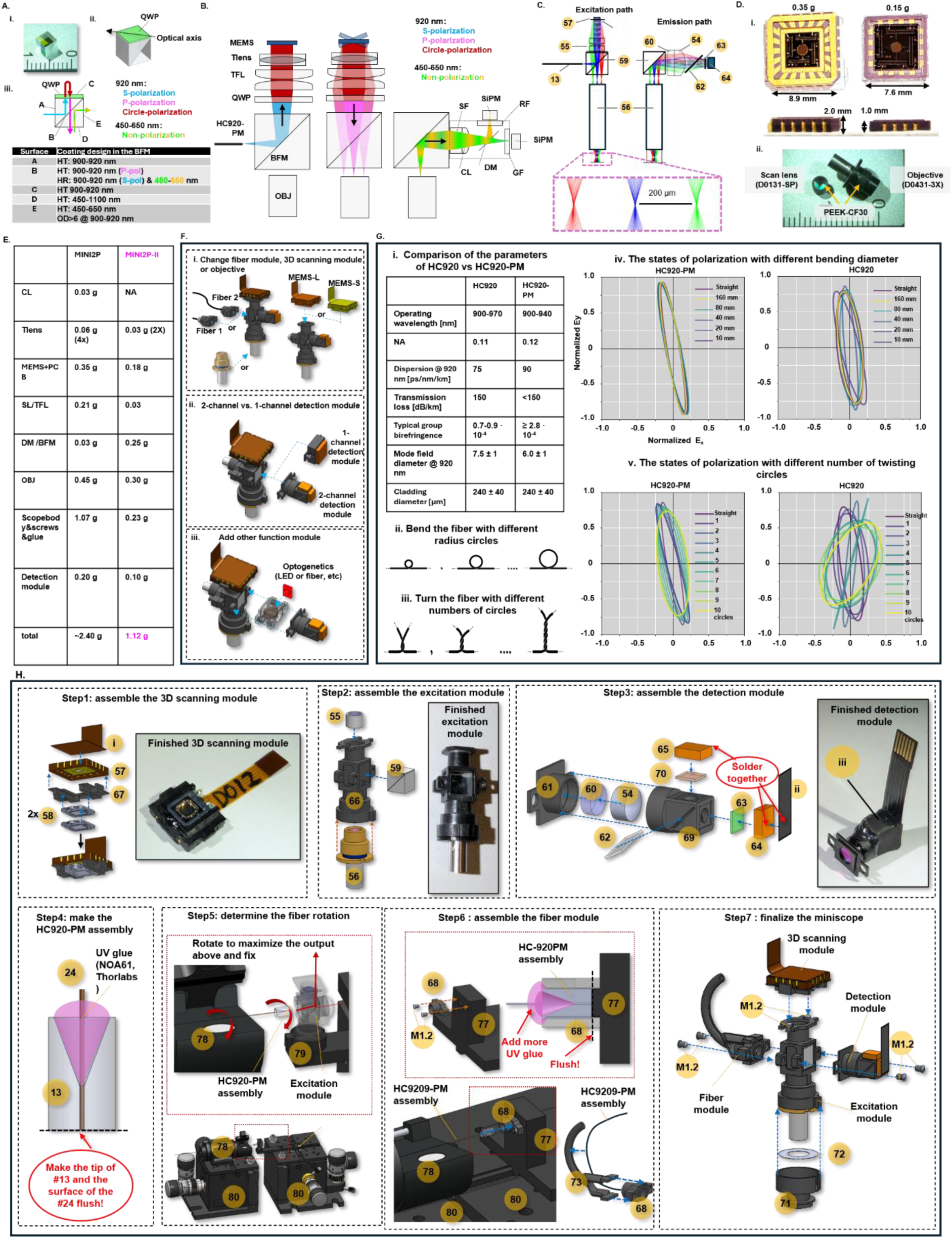
Designing, testing and assembling MINI2P-II, related to Figure 1. A) Beam-folding module (BFM): (i) Photograph. (ii) Mechanical illustration. QWP, quarter-wave plate. The optical axis of the QWP is fixed in the BFM during manufacturing. (iii) Optical surfaces labeled A–E with coating specifications for each surface. B) Optical layout of MINI2P-II showing the static excitation path (fiber to MEMS), steering excitation path (MEMS to objective), and emission path (objective to detector) rendered separately. Wavelengths/polarizations are color-coded: blue, S-polarized 920 nm; purple, P-polarized 920 nm; red, circularly polarized 920 nm; green-yellow, non-polarized 450–650 nm emission. Components: TFL, telecentric f-theta lens; TLENS, tunable lens; MEMS, scanner; OBJ, objective; DM, dichroic mirror; SF, short-pass filter; GF/RF, green/red emission filters; CL, collection lens; SiPM, silicon photomultiplier; HC920-PM, polarization-maintaining hollow-core fiber. C) Optical simulation of the MINI2P-II. (left) Ray tracing from the HC920-PM output to the sample. (right) Ray tracing from the sample to the SiPM detector. Yellow circled numbers correspond to the parts list IDs in the Supplementary. D) Photograph of the lightweight components used in the MINI2P-II. (i) Side-by-side photographs of the MEMS modules used in MINI2P (left) and MINI2P-II (right), illustrating reduced footprint and mass. (ii) Scan lens and objective for MINI2P-II with PEEK-CF30 housings. E) Table of weight comparison for MINI2P vs. MINI2P-II (component-name + mass). F) Illustration of the modularity of MINI2P-II. (i) Fibers, MEMS scanners, and objectives are interchangeable with commercially available options. (ii) Interchangeable 2-channel and 1-channel detection modules (3D models provided on GitHub). Note: the 1-channel module uses a single large-area SiPM (S13360-3075PE, Hamamatsu) that may increase detection efficiency and reduce weight. (iii) Example of adding functional modules, here an optogenetics module (LED or fiber-coupled laser). G) Polarization-maintenance tests comparing HC920-PM with HC920 under controlled bending and twisting. (i) Fiber specifications table. Note the substantially higher group birefringence of HC920-PM, which provides the polarization-maintaining performance. (ii) Bending paradigm: fiber formed into circles of defined diameter. (iii) Twisting paradigm: fiber first bent to a large circle, then rotated to add integer turns. (iv-v) Normalized polarization ellipses measured by a commercial polarimeter (PAX1000IR1/M, Thorlabs) during bending (iv) and twisting (v) for HC920-PM (left) and HC920 (right), illustrating high polarization stability of HC920-PM. H) Step-by-step assembly of MINI2P-II. Full assembly protocol, 2D drawings, 3D models, and materials list will be released on GitHub upon acceptance. Numbers in yellow circles indicate item ID in the parts list in the supplementary document.

**Figure S2.**
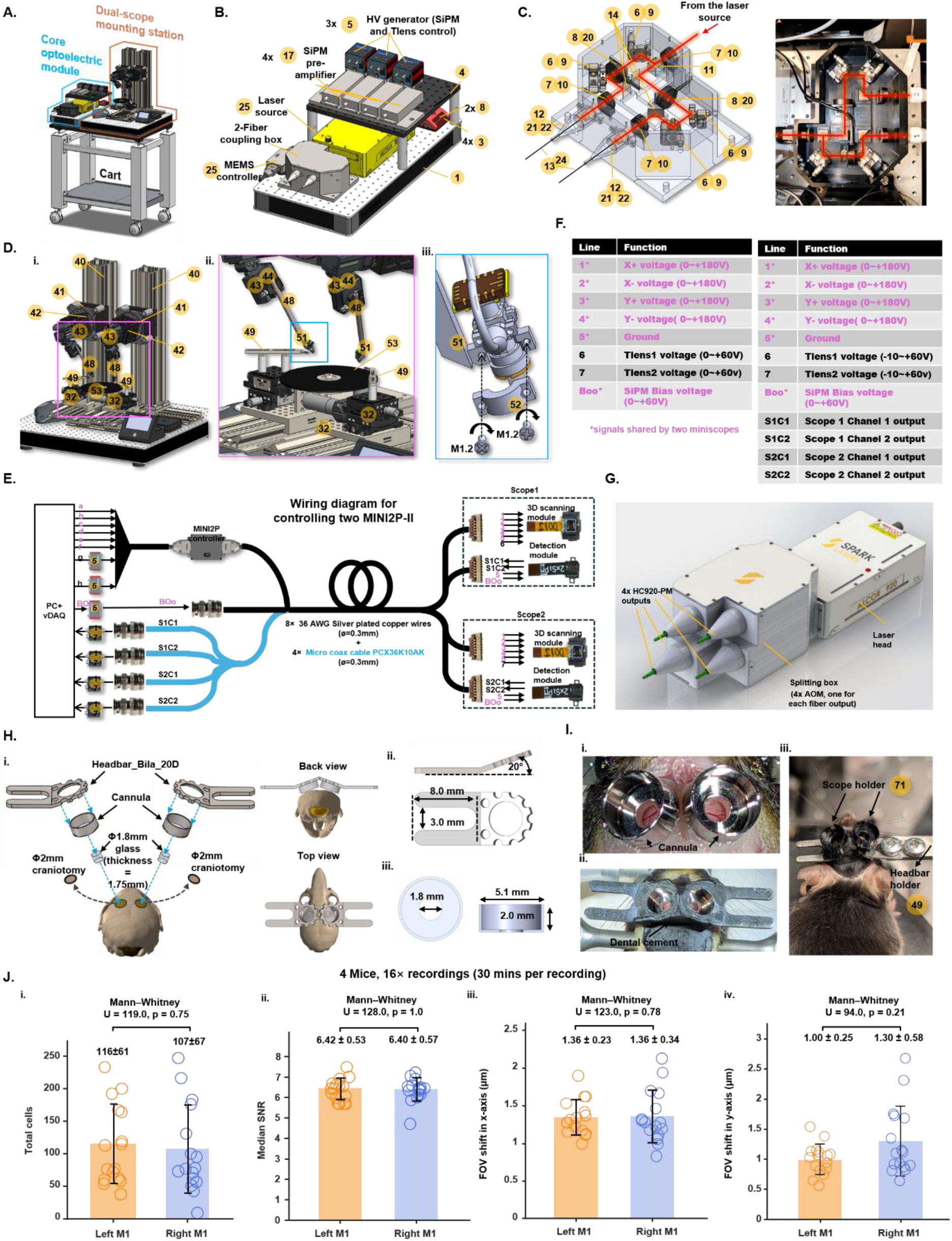
Dual-MINI2P-II system building and surgery protocols, related to Figure 2. A) System overview. Complete dual-scope setup for freely moving recordings. All components fit on a mobile cart < 1 m³ and can be installed in a standard behavior room (no dedicated HVAC/cleanroom required). B) Core opto-electronic module. Key instruments (laser source, MEMS controller, SiPM pre-amplifiers, high-voltage generators for SiPM bias and tunable-lens control and shutter controllers) arranged on a 30 × 60 cm optical breadboard. Yellow circled numbers correspond to **Supplementary parts-list IDs**. C) Beam-splitting and dual-fiber coupling module. Compact module that routes the 920-nm laser to two output fibers (HC920-PM). Updated designs and build tutorials are provided on GitHub. D) Dual-miniscope mounting station. (i) Overview of the head-fixed alignment platform (components on a 60 × 60 cm breadboard). (ii) Close-up of the two miniscope holders and the running wheel used to search for regions of interest (ROIs) during head fixation. (iii) Mechanism for quick fix/release of the MINI2P-II from the scope holder. Updated CAD files are available on GitHub. E) Electrical hardware and cable assembly. Wiring diagram for controlling two miniscopes / four SiPMs (two SiPMs per miniscope) using a 12-wire thin bundle with four 6-pin quick-release connectors at the MEMS PCB and detection modules. The cable combines 8 × 36-AWG silver-plated wires and 4 × micro-coax lines to minimize stiffness and weight. F) Signal definitions. Line-by-line mapping for control/drive and readout signals. Signals marked with an asterisk are shared by both miniscopes. G) Commercial multibeam option. Example of a multi-beam 2P laser providing independent power modulators per fiber output (four outputs shown), enabling per-scope power control. H) Surgery protocol for widely separated targets (e.g. bilateral windows). (i) Diagram of two 2-mm craniotomies (left/right hemispheres) each sealed with a 1.8-mm glass coverslip (thickness 1.75 mm), plus one cannula and one headbar mounted above each window; posterior and dorsal views after implantation shown at the right. (ii) Headbar design. (iii) Cannula design. I) Photographs illustrating the implanted glass windows and cannula (i), mounted headbars with dental cement sealing all gaps (ii), and miniscope baseplates mounted during head-fixed alignment (iii). J) Session-level imaging quality metrics across mice and recordings. Summary across 4 mice and 16 recordings (40 min each). (i) Total cells per recording (Mean±SD). (ii) Median SNR across all recorded neurons for each recording (Mean±SD). (iii) Rigid FOV drift in X/AP (µm; Mean±SD). (iv) Rigid FOV drift in Y/ML (µm; Mean±SD). No significant differences were observed between hemispheres for any metric with Mann–Whitney paired test.

**Figure S3.**
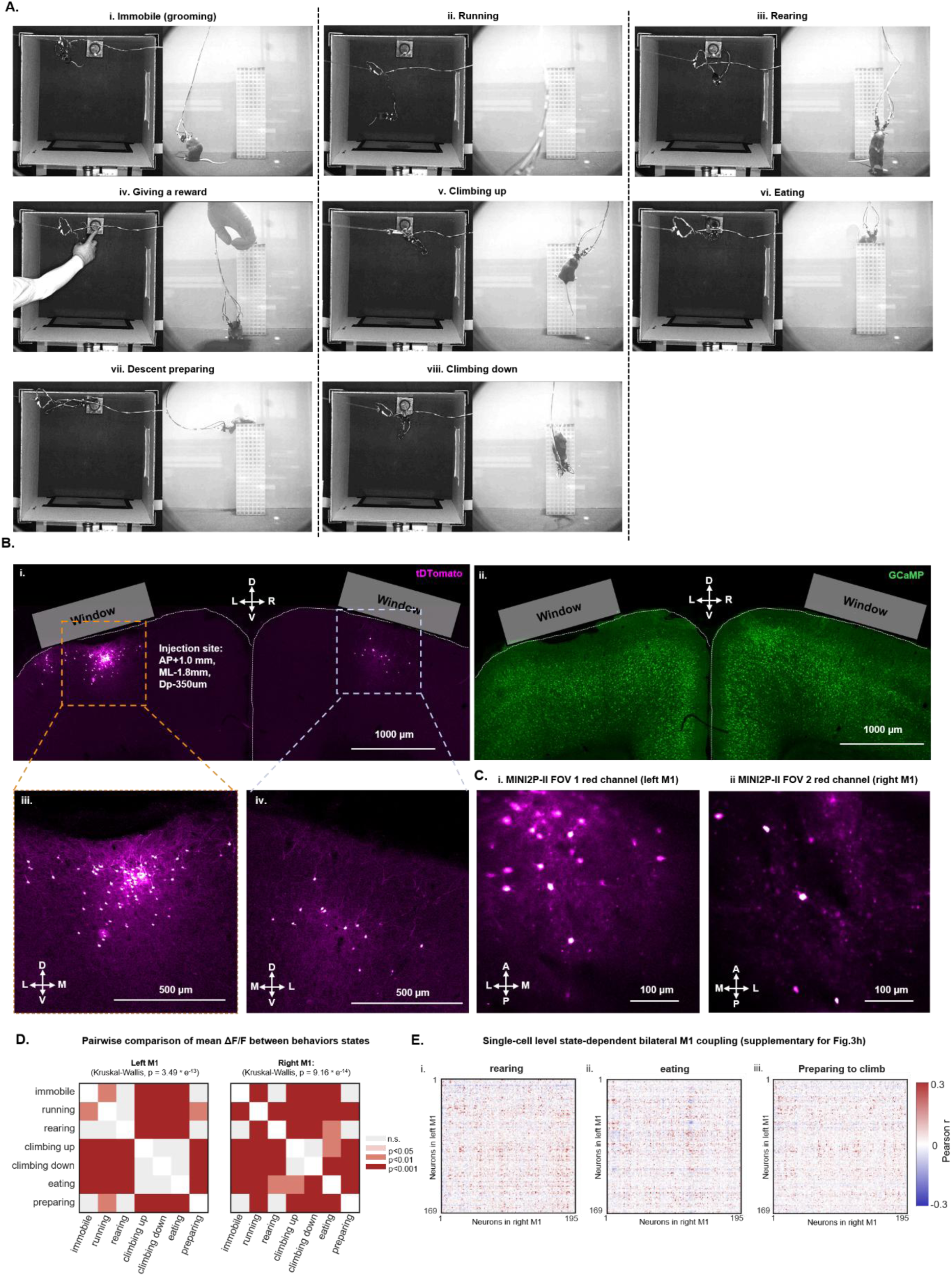
Behavior tracking, histology and supplementary statistics for bilateral M1 imaging during ladder-climbing, related to Figure 3. A) Snapshots from behavioral tracking videos illustrating eight behavioral states. Lef: top-camera view; right: side-camera view. B) Anatomical validation of targeting and labeling. (i) Confocal image (sagittal section) in the red channel showing tdTomato-labeled neurons after retrograde viral injection in left M1 (injection site: AP +1.0 mm, ML −1.8 mm, depth 350 µm). Labeling is spatially restricted and topographic across hemispheres. (ii) Same section in the green channel (GCaMP6s) demonstrating broad, uniform expression. (iii) Zoom of the left M1 injection site illustrating dense local tdTomato labeling. (iv) Zoom of right M1 showing sparse, localized tdTomato-positive neurons projecting to left M1. C) Field-of-view (FOV) selection guided by retrograde label. Red-channel images from MINI2P-II recordings were used to guide baseplating of the MINI2P-II over regions containing tdTomato-labeled neurons in each hemisphere: (i) left M1 (higher density), (ii) right M1 (lower density), consistent with panel a. Axes, A–P and L–R annotated. D) Pairwise statistical comparison of mean ΔF/F across behavioral states for left M1 (i) and right M1 (ii), separately. Each matrix shows bout-level Wilcoxon rank-sum test p-values for all state pairs. Color encodes significance level: light gray, not significant (p ≥ 0.05); faded pink, p < 0.05; medium red, p < 0.01; dark red, p < 0.001. Diagonal cells are white. P-values are printed on each cell. E) Neuron-by-neuron cross-hemispheric Pearson correlation matrices for additional behavioral states not shown in **Fig. 3G**. Same neuron ordering and color scale as in **Fig. 3G**. (i) Rearing. (ii) Eating. (iii) Descent preparation.

**Figure S4.**
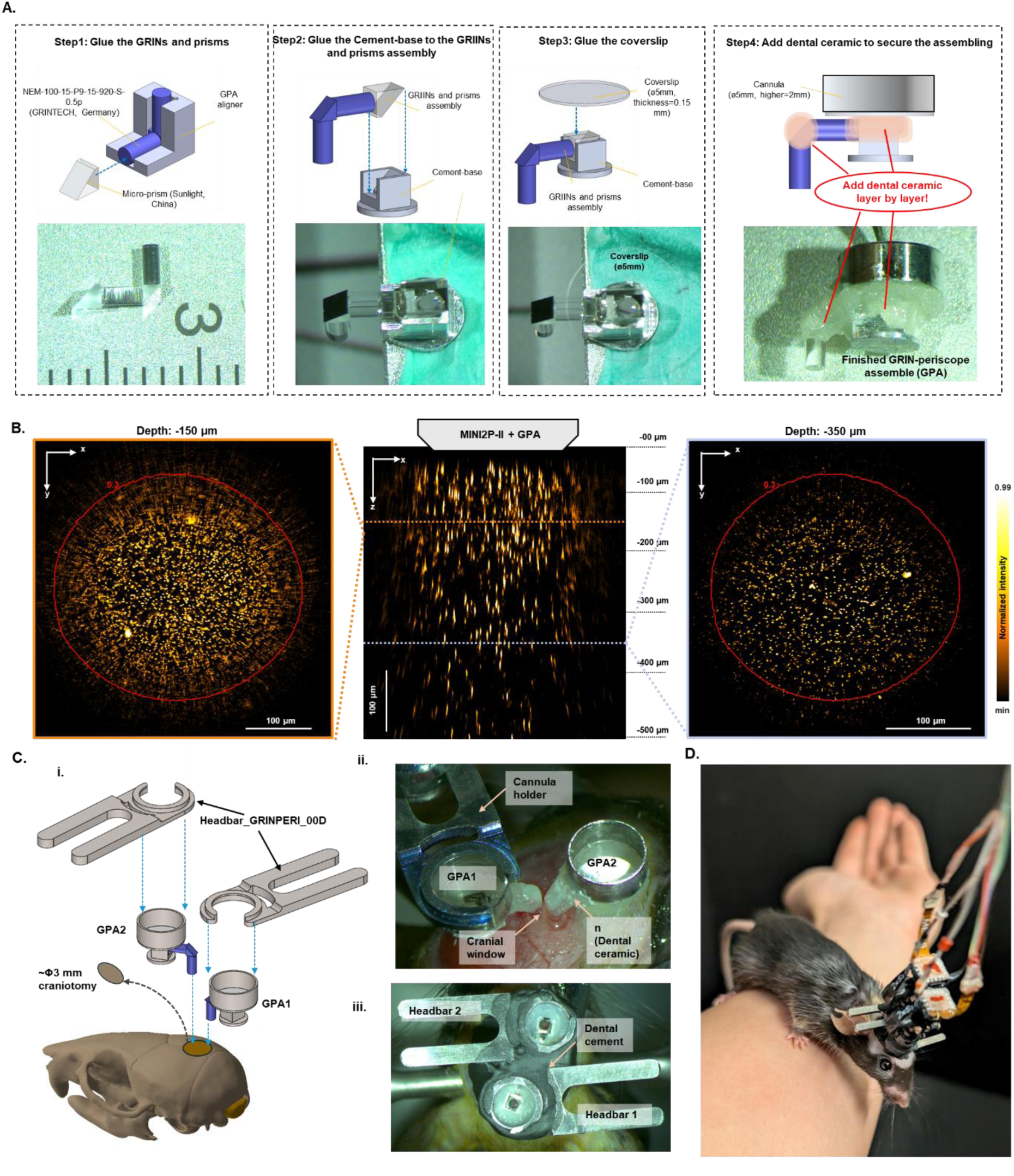
GPA assembling protocol, resolution test, and surgical guidance, related to Figure 4. A) Step-by-step assembly protocol for preparing a GRIN periscope assembly (GPA). Major stages include component cleaning, alignment of GRIN lenses and prisms, bonding with cement base, integration with the coverslip, and securing the assembly with dental ceramic. Complete 2D drawings, 3D models, and a full materials list will be released on GitHub upon paper acceptance. B) Volumetric imaging of 1-µm fluorescent beads through a GPA. Beads were embedded in 1.5% agarose (dilution 1:100). The X–Z projection of the full 500-µm axial range is shown (center). Two example XY planes—150 µm and 350 µm depths—are shown on the left and right, respectively. Red circles indicate the effective FOV (region maintaining ≥30% peak intensity). Note: the effective FOV decreases at deeper planes, and Z-axis tick intervals are non-linear due to depth-dependent magnification (verified by optical simulation in Zemax). C) Surgical guidance for implanting two GPAs targeting adjacent structures within a single cranial window. (i) Diagram of a 3-mm craniotomy accommodating two GPAs, each with its own headbar for independent miniscope alignment. (ii) Intraoperative photograph: GPA2 (right) is fixed to the skull with dental cement covering the cement base and cannula bottom, while GPA1 (left) remains held by the implantation holder. (iii) Both headbars mounted and secured with dental cement, sealing all gaps around the GPA assemblies. D) A freely moving mouse implanted with two GPAs, two headbars, and two MINI2P-II miniscopes.

**Figure S5.**
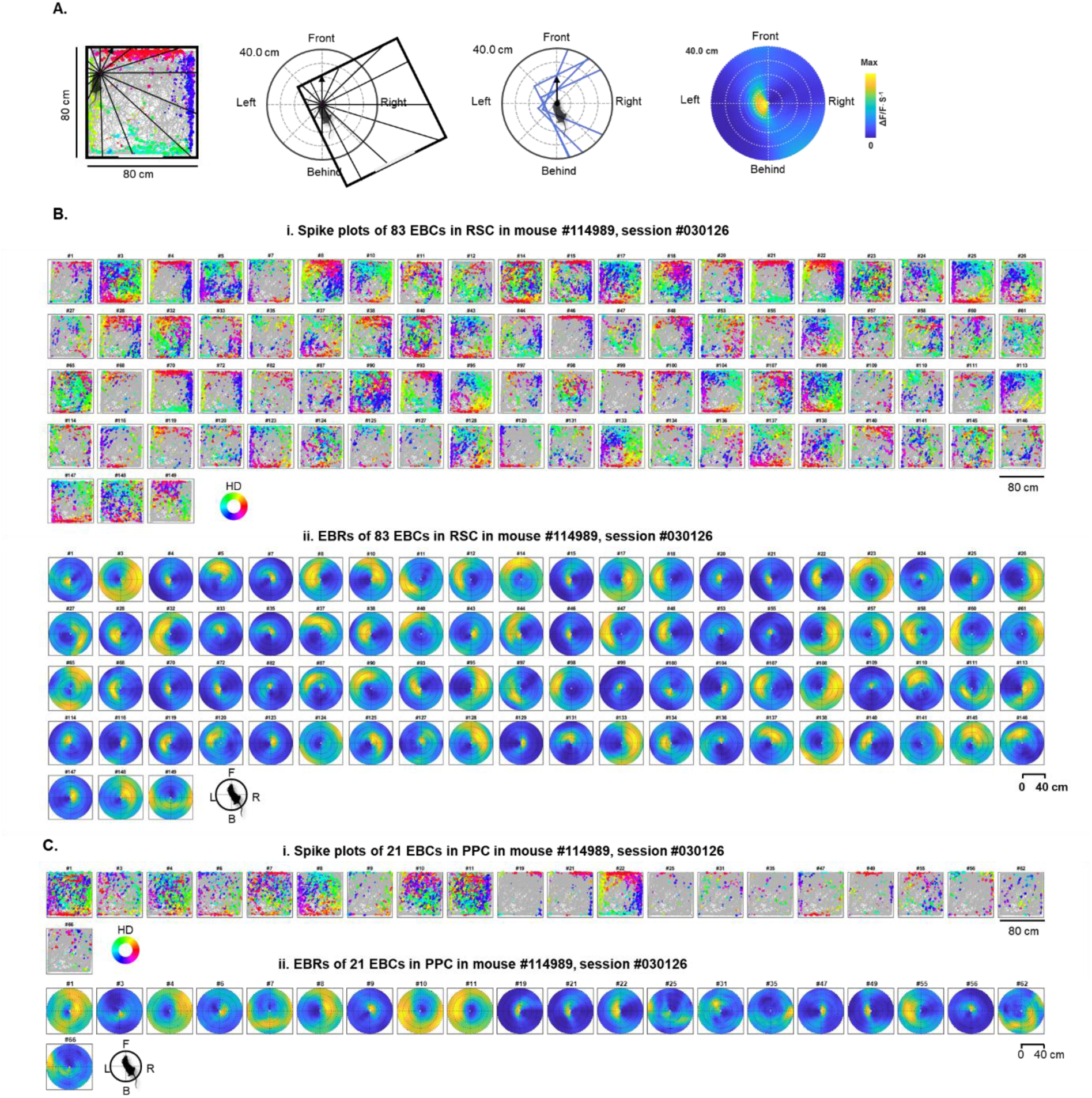
Construction of egocentric boundary rate maps (EBRs) and supplementary EBC datasets, related to Figure 5. A) Algorithm for constructing EBRs, adapted from Alexander et al., 2020. Left and middle panels: for each calcium event (spike), the animal’s head direction (vector arrow) is computed, and the distance to the nearest boundary is determined for all 360° directions (subsample shown for clarity). Middle-left: boundaries within 40 cm are transformed into the egocentric coordinate frame aligned to the current head direction. Middle-right: example egocentric boundary positions for four spikes. Right: resulting egocentric boundary rate map (EBR) for a single neuron. B) All EBCs in RSC for mouse #114989, session #030126 (same session as **Fig. 5A–D**). (i) Spike plots for 83 EBCs, showing the spatial positions of calcium events (trajectory omitted). Events are color-coded by head direction; marker size encodes amplitude (ΔF/F). (ii) Spatial tuning maps for the same 83 cells, each scaled to its peak activity (ΔF/F⋅s⁻¹). C) All EBCs in PPC for mouse #114989, session #030126 (same session as **B**). (i) Spike plots for 21 EBCs, showing spatial positions of calcium events (trajectory omitted), color-coded by head direction as in **B-i**.

**Figure S6.**
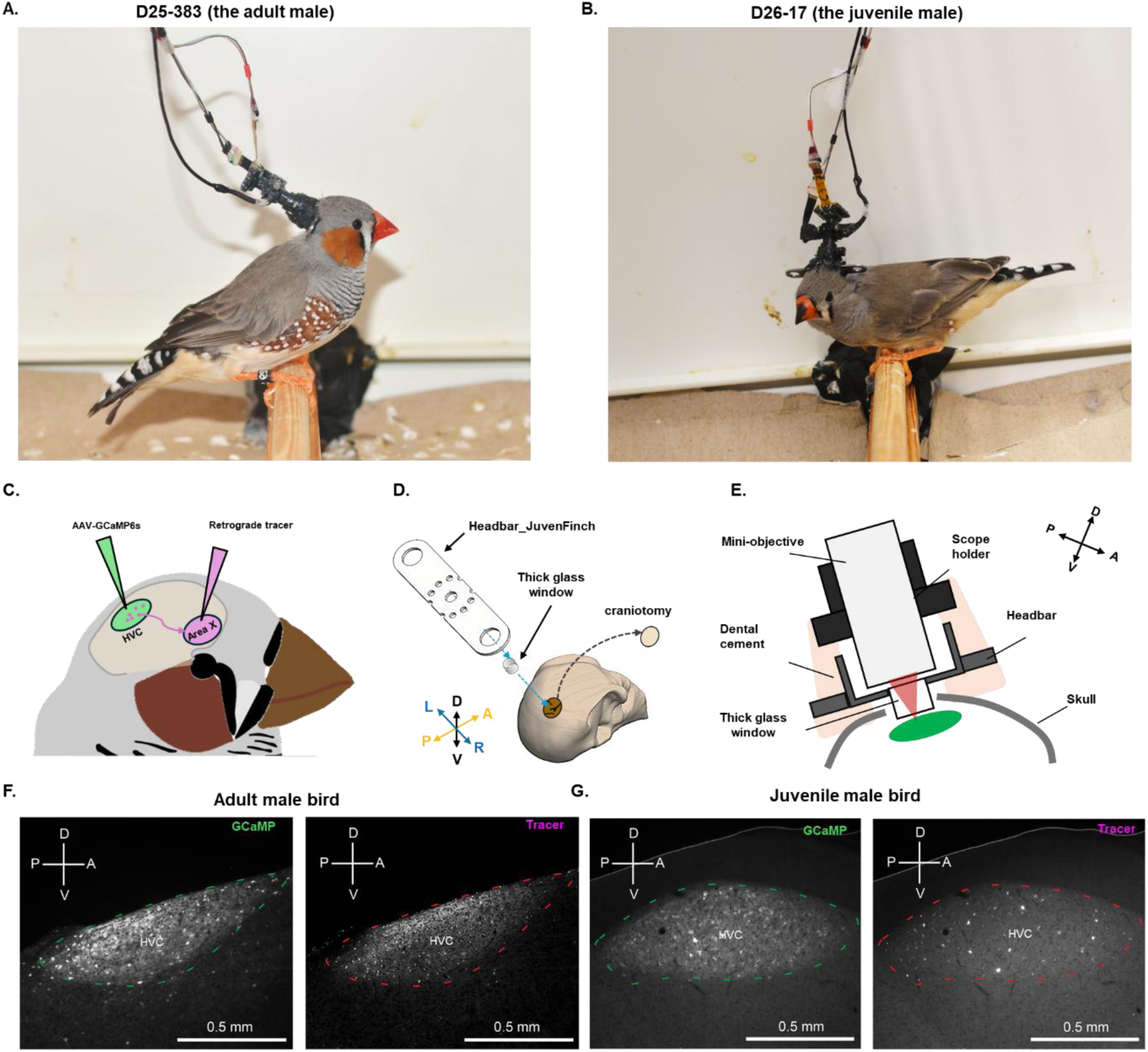
Supplementary information for zebra finch imaging, related to Figure 6. A) Photograph of the adult male bird used in Figure 6 (ID: D25-383) with MINI2P-II mounted on the head. B) Photograph of the juvenile male bird used in Figure 6 (ID: D26-17) with MINI2P-II mounted on the head. C) Viral targeting schematic. AAV–GCaMP6s was injected into HVC. A retrograde tracer was co-injected into Area X, which receives strong projections from HVC (but not from surrounding regions), enabling anatomical disambiguation of recorded areas. D) Surgical guidance for implanting a single cranial window on the bird’s head. The headbar (“Headbar_JuvenFinch”) was designed to achieve low weight, high stability, and maximal flexibility to accommodate different skull sizes in adult and juvenile birds. E) Schematic of the MINI2P-II miniscope mounted on top of the cranial window. F) Representative histological verification showing that GCaMP6s expression (left) and retrograde labeling (right) are restricted to HVC of an adult male zebra finch. Dashed lines in the images indicate the border of HVC in the sagittal sections. G) Representative histological verification showing GCaMP6s expression (left) and retrograde labeling (right) restricted to HVC in a juvenile male zebra finch (sagittal sections). Dashed lines as in F.

**Video S1. Dual-color MINI2P-II imaging in visual cortex during unrestrained movement**, related to Figure 1. Top left: GCaMP6s-expressing neurons (green channel). Top right: tdTomato-labeled astrocytes (red channel). Bottom right: overhead tracking of mouse behavior. The video plays at 3× speed, and a 3-frame running average is applied to both 2P imaging channels for visualization.

**Video S2. Bilateral M1 imaging during ladder-climbing behavior**, related to Figure 2. Top left: schematic of the two recording sites in left and right primary motor cortex (M1). Top middle: MINI2P-II imaging from left M1. Top right: MINI2P-II imaging from right M1. Bottom left: overhead behavioral tracking. Bottom right: side-view behavioral tracking. The video plays at 3× speed, with a 3-frame running average applied to both FOVs for visualization.

**Video S3. Simultaneous ipsilateral RSC–PPC imaging during natural behavior**, related to Figure 4. Top left: MINI2P-II imaging of retrosplenial cortex (RSC) in the right hemisphere. Top right: MINI2P-II imaging of posterior parietal cortex (PPC) in the same hemisphere. Bottom left: schematic of the two GPA-shifted recording sites. Bottom right: overhead tracking of mouse behavior. The video plays at 3× speed, with a 3-frame running average applied to both FOVs for visualization.

**Video S4. 2P calcium imaging during adult male zebra finch courtship singing**, related to Figure 6. Top left: MINI2P-II imaging of HVC in a freely moving adult male zebra finch; a white indicator box appears when the bird is singing. Top right: wide-view behavioral monitoring from an external camera. Bottom right: synchronized spectrogram extracted from the audio track (color scale: amplitude from ∼10 Hz to 10 kHz). A moving vertical bar marks the current time point corresponding to the 2P imaging and video frame. The video plays in real time, with a 3-frame running average applied to the 2P imaging for visualization.

**Video S5. 2P calcium imaging during social interaction in a juvenile zebra finch**, related to Figure 6. Top left: MINI2P-II imaging in a juvenile finch. Top right: behavioral monitoring from an external camera. Bottom right: synchronized spectrogram extracted from the audio track (color scale: amplitude from ∼10 Hz to 10 kHz). A moving vertical bar marks the current time point synchronized with 2P imaging and video frames. The video plays in real time, with a 7-frame running average applied to the 2P imaging for visualization.

## EXPERIMENTAL MODEL DETAILS

### Animals

Mice experiments were performed according to the Norwegian Animal Welfare Act and the European Convention for the Protection of Vertebrate Animals used for Experimental and Other Scientific Purposes, permit numbers 18013, 6021 and 7163. Mice were housed in a specific-pathogen-free (SPF) facility following recommendations from the Federation of European Laboratory Animal Science Associations (FELASA; Mahler et al., 2014). We used transgenic mice expressing GCaMP6s under the CaMKII promoter (Camk2a-tTA; tetO-G6s; Wekselblatt et al., 2016). Visual cortex (VC) recordings in **Figure 1** were obtained from one female mouse. Bilateral M1 recordings (**Figures 2**, **S2**, **3**, and **S3**) were collected from four transgenic mice. Recordings in retrosplenial cortex (RSC) and posterior parietal cortex (PPC) (**Figures 4**, **S4**, **5**, and **S5**) were obtained from an additional cohort of three transgenic mice. All animals were 12–24 weeks old at the time of surgery and 16–28 weeks old at the start of behavioral pre-training (see **Supplementary Document Part 2: imaging conditions** for more details). Before implantation, mice were group-housed with up to three littermates (cage size: 32 × 17 × 15 cm). Mice were handled daily and pre-trained in the recording arena until they were fully habituated and foraged freely. After implantation, mice were housed individually in transparent Plexiglass cages (36 × 24 × 26 cm). All mice were maintained on a 12 h light/12 h dark cycle and tested during the dark phase. Animals were never subjected to food or water restrictions. Health status was monitored daily throughout the experimental period.

Bird experiments were performed in accordance with the ethical principles and guidelines for animal experiments in Germany, and all experimental protocols were approved by the government of Upper Bavaria (ROB-55.2-2532.Vet_02-24-114). Young birds aged between 25 and 60 days and adult birds aged between 130 and 365 days were obtained from the breeding colony in Seewiesen, Germany. All birds received injections of a GCaMP AAV virus into HVC and with Red Retrobeads (Lumafluor Inc. NC, USA) into Area X. Young birds were 25-30 days old at the time of surgery. Before implantation, birds were housed in an aviary (1 × 2 × 2 m). All birds were maintained on 15 h light/9 h dark cycle. Animals were never subjected to food or water restrictions. Health status was monitored daily throughout the experimental period.

## METHOD DETAILS

### Bilateral M1 surgery

Anesthesia was induced by placing the mouse in a sealed glass chamber filled with 5% isoflurane delivered at 1.2 L/min. Animals were then transferred rapidly to a stereotaxic frame equipped with an isoflurane nose cone, where anesthesia was maintained with 0.5–2% isoflurane in air (1.2 L/min), adjusted based on physiological monitoring. Two systemic analgesics (Temgesic and Metacam) were administered subcutaneously, and a local anesthetic (Marcain) was injected subcutaneously over the planned craniotomy region. Mannitol (150 mg/ml) was injected intraperitoneally 30–60 min before the durotomy to reduce intracranial pressure. The scalp was resected to expose the entire dorsal skull, and the periosteum was removed while leaving the bone intact. Two circular craniotomies (diameter slightly >1.8 mm) were drilled bilaterally over M1 (centered 2.0 mm lateral and 1.0 mm anterior to bregma). The dura remained intact throughout the procedure. To enhance adhesion, a thin coat of Optibond (Kerr, CA, USA) was applied to the skull followed by a thicker layer of Charisma (Kulzer, Hanau, Germany), both cured under UV light.

A customized imaging assembly—consisting of a 1.8-mm-diameter coverglass (Sunlight, Fuzhou, China), a titanium cannula, and a titanium headbar—was pre-glued using optical glue (NOA61, Norland, NJ, USA). The assembly was inserted into each craniotomy using a custom headbar holder mounted on a stereotactic micromanipulator (1760, Kopf, CA, USA). The holder was tilted 20° laterally to ensure that the coverglass rested approximately parallel to the dorsal surface of M1. The assembly was lowered until the coverglass gently contacted the brain surface. The interface between the headbar and the skull was sealed first with Venus (Kulzer, Hanau, Germany), and remaining gaps were then filled with dental cement (Metabond, Parkell, NY, USA). After curing, the headbar was released from the holder. The chronic windows were finally protected with a layer of Kwik-Cast (World Precision Instruments, FL, USA).

During the same surgical session, a retrogradely transported AAV was injected into left M1 to label local layer 2/3 neurons and callosally projecting neurons from the contralateral M1 (Tervo et al., 2016). After exposing the left cranial window, a glass capillary (Nanoject III) loaded with rAAV2-CAG-tdTomato (Addgene plasmid #59462; RRID: Addgene_59462, diluted 1:20) was lowered to target layer 2 of the M1 forelimb subregion (1.8 mm lateral, 1.0 mm anterior to bregma; 0.35 mm depth). A total of 50 nL was injected at 30 nL/min. The pipette remained in place for 10 min before withdrawal to minimize backflow.

### Assembling the GRIN periscope assemble (GPA)

The GRIN periscope assembly (GPA) consisted of a custom titanium cannula (5 mm diameter, 1 mm depth), a 5-mm-diameter coverglass (CS-5R, Warner Instruments, MA, USA), a 1.3 × 1.3 × 1.6 mm³ custom prism (Sunlight, Fuzhou, China), a pre-mounted GRIN lens module (NEM-100-15-P9-15-920-S-0.5p, GRINTECH, Germany), and an aluminum cement base designed to ensure secure fixation to the skull using dental cement (see **Fig. S4C-ii, iii**). These components were aligned and bonded using optical glue (NOA61, Norland, NJ, USA; **Fig. S4A**). After alignment, the optical glue was cured with UV light (CS20K2, Thorlabs, NJ, USA). To prevent stress-induced cracking during implantation, multiple very thin layers of dental ceramic (Charisma, Kulzer, Germany) were applied at all connection interfaces, for example, between the prism and the coverglass and between the prism and the GRIN lens (**Fig. S4A**, final step). The new glue was applied after the previous glue was UV-cured.

### RSC and PPC surgery

Anesthesia was induced by placing each mouse in a sealed glass chamber filled with 5% isoflurane at 1.2 L/min, after which the animal was transferred rapidly to a stereotaxic frame equipped with an isoflurane nose cone. Anesthesia, airflow, and analgesic regimens were identical to those used in the M1 surgeries. Mannitol (150 mg/ml) was administered intraperitoneally 30–60 min before the durotomy to reduce intracranial pressure. The scalp was removed to expose the full dorsal skull, and the periosteum was cleared while leaving the bone intact. A circular craniotomy (≈3 mm diameter) was made over the right hemisphere, centered 1.5 mm lateral and 2.5 mm posterior to bregma leaving the dura intact. The dura remained intact. The pitch of the head was adjusted so that the anterior and posterior edges of the cranial window were parallel to the horizontal plane.

The first GRIN periscope assembly (GPA) was mounted onto a custom cannula holder connected to a stereotaxic micromanipulator (1760, Kopf, CA, USA; **Fig. S4C-ii**). For implantation over RSC, the GPA was lowered slowly until the bottom of the lower GRIN lens made gentle contact with the brain surface and the left edge of the lens was positioned adjacent to the right border of the sagittal sinus. The gap between the GPA’s cement base and the skull was filled with dental ceramic (Charisma, Kulzer, Hanau, Germany), and the GPA was then released from the holder.

The second GPA was subsequently loaded onto the same holder and positioned over PPC. It was lowered until the lower lens contacted the brain and the center of the lens aligned with PPC coordinates (AP −2.0 mm, ML +2.0 mm relative to bregma). As with the first implant, the gap beneath the cement base was filled with dental ceramic before the GPA was released.

All remaining gaps between the GPAs and the skull were sealed with Venus (Kulzer, Hanau, Germany). Next, a custom headbar was mounted on top of each GPA (**Fig. S4C-iii**), and all remaining spaces between the headbars and the skull were filled with dental cement (Metabond, Parkell, NY, USA). After implantation, the chronic cranial window was protected with a layer of Kwik-Cast (World Precision Instruments, FL, USA).

### Zebra-finch surgery

Birds were given an injection of metamizole (100 - 150 µg/g body weight) into the pectoral muscle 15 minutes before surgery. They were then anesthetized with isoflurane inhalation 0.8 - 1.8% at 0.5 l O2/min). After administration of the local anesthetic procaine (2%), a vertical incision was made in the skin over the skull and a craniotomy was performed. For Retrobeads injection into Area X of adult (or juvenile) birds, craniotomy was placed at 1.4 (1.4) mm lateral, 5.6 (5.5) mm rostral and 3.3 (3.3) mm ventral relative to the Y-sinus zero point, with an 80-degree angle between the ear bar and the beak. 100 nl (100%) of rhodamine-labeled Retrobeads (Lumafluor Inc. NC, USA) was injected into Area X. For GCaMP injection into HVC, the craniotomy was placed at 2.4 (or 2.7 for juvenile) mm lateral, 0 (-0.15 for juvenile) mm rostral and 0.3 (0.35 for juvenile) mm ventral related to the Y-sinus zero point, with a 45-degree angle between the ear bar and the beak. 400 nl of viral vector (0.3-3x10^13^ vg/ml) was injected using glass pipettes attached to a manually operated hydraulic injector. To avoid backflow during injection, pipettes were left in place for 5 minutes before withdrawal.

In adult birds, a chronic cranial window with a headpiece was placed over the HVC immediately after viral vector injection. In young birds, this was done 10 days after viral vector injection. The craniotomy involved removing the skull above the HVC while leaving the dura intact to allow for long-term imaging. To enhance adhesion and harden the skull of young birds, a thin layer of Optibond (Kerr, CA, USA) was applied to the skull followed by a thicker layer of Charisma (Kulzer, Hanau, Germany), both cured using UV light. A customized glass plug (1.8 mm diameter and 1.5 mm long, Sunlight, Fuzhou, China) was fitted into the cranial opening and secured to the skull with Kwik Sil (World Precision Instruments, FL, USA) and UV-curing dental cement (Venus, Kulzer, Hanau, Germany). A custom-made titanium headpiece with a 5-mm diameter opening were bonded to the skull with dental cement (Paladur, Kulzer, Hanau, Germany). Finally, a 5 mm diameter coverglass (Sunlight, Fuzhou, China) was glued to the glass plug using optical glue (NOA61, Norland, NJ, USA). After anesthesia recovery, birds were administered post-treatment meloxicam (0.2 mg/kg body weight) injected into the pectoral muscle for analgesia.

### Histology

At the end of all mouse experiments, the animals were anesthetized with isoflurane and euthanized with an overdose of pentobarbital. Transcardial perfusion was performed first with 0.9% physiological saline, followed by 4% formalin or paraformaldehyde (PFA). The implantation sites were carefully dissected with a small portion of the surrounding skull left intact to preserve the chronic cranial window and maintain the relative alignment between the cortex and cerebellum during fixation.

Brains were post-fixed in 4% formalin/PFA for 24 hours at 4°C. Following fixation, the brains were gently separated from the skull while preserving the connection between cortex and cerebellum. Brains were decanted from formalin/PFA and cryoprotected with 30% sucrose for at least 48 hours at 4°C. Whole brains were subsequently embedded in OCT mounting media (VWR international). Tissue was then sectioned in the coronal plane at 40 μm sections with a cryostat (Thermo Scientific CryoStar NX70). For GFP staining, free-floating sections were washed in PBS 2x10 min. The sections were subsequentially submerged in blocking solution containing PBS, 3% bovine serum albumin (BSA) and 0.3% Triton X-100 for 60 min. Sections were incubated overnight at RT on a shaker in wells containing rabbit anti-GFP primary antibody (Invitrogen; catalog #A-11122; RRID:AB_221569; dilution 1:1000) prepared in PBS with 3% BSA and 0.3% Triton X-100. The following day, slices were washed 3x10 min in PBS containing 1% BSA and 0.1% Triton X-100 and incubated for 90 min RT on a shaker in wells containing Donkey anti-Rabbit IgG (H+L) Alexa Fluor 488 secondary antibody (Invitrogen; catalog #A-21206; RRID:AB_2535792; dilution 1:750). Sections were then washed in PBS containing 1% BSA and 0.1% Triton X-100 1x10 min and in PBS 2x10 min at room temperature and mounted with ProLong™ Gold Antifade Mountant (Invitrogen; catalog #P36930) on Superfrost Plus Adhesion Microscope Slides (Epredia, J1800AMNZ). The positions of the glass window and the resulting fluorescent were imaged using a Zeiss laser scanning microscope 990 and Zen (black edition) software (Carl Zeiss, Jena, Germany).

Birds were euthanized with an overdose of isoflurane, then perfused with pH 7.4 phosphate buffered saline (PBS) followed by 4% paraformaldehyde in PBS (PFA). The brain was removed and fixed in 4% PFA for 5-7 days followed by 10% and 30% PBS-sucrose for 1 day each. Brain sections (20 µm) were produced with a freezing microtome (CM1325, Leica, Wetzlar, Germany). Sections were embedded with antifade mounting medium (VEC-H-1000, BIOZOL, Hamburg, Germany) on glass slides and imaged with a wide field microscope (EVOS, Thermo Fischer Scientific, CA, USA, objective: Olympus, 20x0.8 air, SN: N5702000) and a confocal microscope (SP5, Leica, Wetzlar, Germany, objective: Leica, 40x1.25 oil, SN: 506252).

### Open-field task

Recordings began after full recovery from surgery. Before experiments, mice were gradually habituated to handling, the recording setup, and the behavioral task. On each experimental day, mice were placed in the recording room during final preparations to promote familiarity with the environment. All behavioral sessions were conducted in a dark room illuminated only by a ceiling-mounted warm-white LED strip to encourage exploration. A session consisted of a single recording lasting either 29 min (behavior test) or 40 min (RSC/PPC experiments). Each session consisted of a single continuous recording lasting either 29 min (behavioral experiments, **Figs. 2D** to **2J**) or 40 min (RSC/PPC experiments, **Figs. 4** and **5**). During each session, animals freely explored an 80 × 80 cm² square box. A single sheet of laminated white A4 paper was affixed to one wall to provide a polarizing cue. Mice were placed in the arena and allowed to move freely while the experimenter periodically scattered small cookie crumbs to maintain exploratory behavior. Animal position was tracked from above using a monochrome NIR camera (CS505MU1, Thorlabs, NJ, USA) equipped with a 12-mm lens (F1.4, Navitar). The arena and floor were uniformly illuminated by 8–12 LED arrays emitting 850-nm infrared light positioned at fixed locations around the arena. An 875-nm short-pass filter (#86-114, Edmund Optics, USA) was mounted in front of the camera to block any residual 920-nm light leakage from the miniature 2-photon microscope. Tracking videos were acquired using the manufacturer’s software (ThorCam, Thorlabs, NJ, USA) in external-trigger mode. At the start of each 2-photon imaging frame, a TTL pulse (0–5 V) generated by the vDAQ card (see STAR Methods, “MINI2P-II system”) triggered the capture of a single video frame by the tracking camera. This ensured frame-accurate synchronization between animal tracking and 2-photon imaging at all times.

### Locomotion behavioral testing

To evaluate the impact of mounting two miniscopes and their connection cables on naturalistic behavior, we used the same four transgenic mice prepared for bilateral M1 imaging. The aim of these experiments was specifically to assess the combined effect of *dual miniscopes plus cables* on free behavior, rather than isolating the effects of miniscope weight or cables independently. In our previous work (Zong et al., Cell 2022), we demonstrated that (i) a single miniscope weighing ≤3 g does not disrupt naturalistic foraging behavior, and (ii) cable stiffness has a large impact on movement than miniscope weight alone. Therefore, experiments with only one miniscope or with an unconnected miniscope were not repeated here. Each mouse participated in two “with” sessions (two miniscopes + cables attached) and two “without” sessions (no miniscopes or cables). Session order was interleaved across days (e.g., Day 1: without; Day 2: with; Day 3: without; Day 4: with) to minimize confounds from day-to-day behavioral variability. Prior to testing, animals were habituated to the weight of the miniscopes using 3D-printed dummy attachments mounted on the implanted headbars for at least 5 days. At the start of each session, mice were head-fixed briefly while dummies were removed. We then either mounted the real miniscopes and connected cables (“with” sessions) or attached nothing (“without” sessions). Mice were then gently placed into the open-field arena and performed a 30-min free-foraging task using the same behavioral setup described above. The first 1 min of each session was excluded from analysis to remove transient behavioral adaptation to entering the dark arena. After each session, mice were retrieved using a familiar transfer tube, the dummy scopes were remounted, and animals returned to their home cages. All testing occurred at similar times each day, and all four animals were run on each test day to minimize circadian influences on activity levels.

### Behavioral Quantification

Six behavioral metrics were analyzed across “with” and “without” sessions:

1. Accumulated running distance over time
2. Total running distance
3. Distance traveled in the center of the arena
4. Mean running speed
5. Median running speed
6. Momentary tortuosity of the running path

To compute these measures, x–y head positions, *X*(*t_i_*)and *Y*(*t_i_*), were extracted from the tracking pipeline (see Methods: Tracking Data Processing) and smoothed with a 0.5-s moving average. Here, *t_i_*denotes the *i*th video frame (frame interval Δ*T* = 0.07s). The inter-frame running distance was computed as:

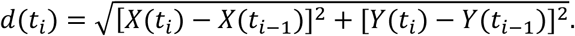

The total running distance was the sum of all inter-frame distances over the session. The distance traveled in the arena center was defined as the total distance traveled within ±15 cm in both x and y directions relative to the arena center. Momentary running speed was defined as:

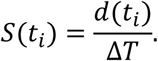

Mean and median speed values were computed over the entire recording. Momentary tortuosity was defined as in Zong et al. (Cell 2022). For each time point, a ±1.25-s sliding window centered at time *t* was used. The path length within the window was:

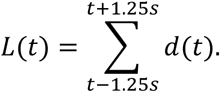

The straight-line distance between window endpoints was:

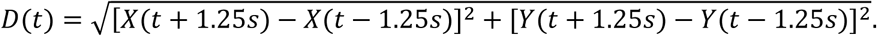

Momentary tortuosity was then:

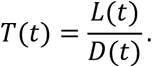

Tortuosity was evaluated only during running periods (*S* > 2.5 cm/s).

### Reward-motivated ladder-climbing paradigm

This experiment was performed in four male transgenic mice (Camk2a-tTA; tetO-G6s) implanted with two 1.8-mm glass windows over left and right M1 and recorded simultaneously with two MINI2P-II miniscopes. Calcium activity was imaged while mice climbed up and climbed or jumped down from a tall platform placed inside a 50-cm-wide square box with a black rubber floor and 50-cm-high black acrylic walls. A white polylactic acid (PLA) platform (22 cm height; 7 × 7 cm top area) was 3D-printed with one vertical face (22 × 7 cm) containing an array of square holes (6 × 6 mm²) spaced 4 mm apart horizontally and vertically, serving as a ladder for climbing. A 1-kg steel block was mounted inside the platform to ensure stability. A lateral monochrome camera (acA2040-90umNIR, Basler, Ahrensburg, Germany) equipped with a 3.5-mm lens (#89-410, f = 3.5 mm, Edmund Optics, NJ, USA) was positioned outside the arena, filming through a hole in the wall directly facing the ladder surface. The same overhead NIR tracking camera used in the open-field task was used simultaneously for top-down position tracking. The two cameras were hardware-synchronized.

After habituation to the miniscope weight, the assay was performed daily for 5 consecutive days, starting on Day 1 with trials in which the MINI2P-II miniscopes were already mounted. The platform was placed inside the arena before each trial, and a highly palatable reward (a small amount of sweet cream) was placed on the top surface. The platform was positioned near the wall opposite the lateral camera, with the ladder face oriented toward the camera. Mice were trained to climb vertically to reach the reward and then to either climb or jump down from the platform before initiating another ascent. After the mouse descended, the next reward was delivered manually, and a new reward was placed on top of the platform. Each session lasted up to 40 min or ended early if the mouse did not climb for more than 5 min.

Seven behavioral states were annotated manually by visual inspection of synchronized video recordings. The climbing-up phase began when all four paws left the ground and ended when all four paws were on the top platform. The climbing/jumping-down phase starts after a preparation period in which the rear paws grasped the platform edge, followed by the descent or fall interval, which lasted from the moment all four paws left the platform until the rear paws first contacted the floor.

### Tracking zebra finch singing behavior

For song recording, birds were housed temporarily in a custom-made soundproof chamber (120 x 50 x 50 cm) for maximum 5 days. Each chamber was equipped with a microphone (TC20, Earthworks, NH, USA). The sounds were recorded digitally with a sampling rate of 44.1 kHz and an accuracy of 16 bits using SAP2011 software (Tchernichovski et. al., 2000). To monitor the vocal developments of young birds and to assure that each bird reached the stage of full-songs, young birds were housed multiple times in soundproof chambers.

### MINI2P-II miniscope

Linearly-polarized laser light enters the miniscope through the HC920-PM fiber and enters the beam-folding module (BFM; Sunlight, Fuzhou, China; **Fig. S1A**). The BFM is described in detail in **Fig. S1A**; briefly, it routes light differently depending on input direction, wavelength, and polarization, enabling a highly compact optical path. The S-polarized beam exiting the fiber is directed upward through a pre-aligned quarter-wave plate (QWP; fast axis oriented at 45°) mounted on the BFM, and then into a telecentric f-theta lens (TFL; D0131-SP, Domilight, Nanjing, China), which collimates the beam. The collimated light passes through two tunable micro-lenses (Tlenses; PIF.15.P02, Polight, Horten, Norway) that allow voltage-controlled axial focus modulation for imaging at different depths in the brain. From the Tlenses, the beam enters a 2-axis MEMS scanner (A3I12.2-1200AL-LCC16-TW, Mirrorcle, CA, USA) that controls the angular deflection of the excitation beam. The reflected light then passes back through the Tlenses and TFL. When traversed in this reverse direction, the TFL converts angular deflection into x–y translation at the sample plane, and changes in beam convergence into axial shifts in focus. The TFL’s flat-field design maintains a uniform focal plane and minimal spot distortion across scanning angles. After leaving the TFL, the beam re-enters the BFM. Because the beam has passed through the QWP twice in opposite directions, its polarization has rotated by 90°, enabling it to be transmitted through the BFM toward the objective lens (**Fig. S1B**). The objective (D0431-3X, Domilight) uses standard Thorlabs M5-0.5 threading and maintains the same optical distance to the imaging plane as other 2P miniscope objectives, making them interchangeable. Fluorescence emitted from the sample is collected by the same objective and is directed into the BFM, which reflects it toward an absorptive low-pass filter (BG40-920-D2.8-L1.0, Sunlight) and a collection lens (49-168, Edmund Optics, NJ, USA). The collection lens focuses short-wavelength (500–550 nm) and long-wavelength (595–665 nm) emission components onto two onboard silicon photomultipliers (SiPMs; S13360-1375PE, Hamamatsu, Japan). Before reaching the detectors, fluorescence is split by a custom dichroic mirror (Sunlight; ID #62 in **Figs. S1C, S1H** and the **Supplementary Document Part 3: parts list**) and filtered by two additional custom bandpass filters (IDs #63 and #70).

The MEMS scanner and the two Tlenses are assembled together as a 3D scanning module consists of (Step 1 in **Fig. S1H**). The excitation module includes the TFL and BFM, which are glued directly to the scope body with the objective screwed into place (Step 2). The 2-channel detection module comprises the low-pass filter, collection lens, second dichroic mirror, color filters, and two SiPMs assembled into a compact unit (Step 3). The HC920-PM fiber assembly, the 3D scanning module, and the detection module are mounted onto the excitation module and secured with two M1.2 screws. Electrical connectors for the scanning and detection modules are designed for easy plug-in removal. This high-modularity architecture facilitates rapid repair, component upgrades, and integration of additional functional modules (**Fig. S1F**).

The compact, tightly matched design simplifies optical alignment and mechanical mounting. Complete assembly of a single MINI2P-II microscope consists of seven discrete steps and requires approximately one hour (**Fig. S1H**).

### Beam splitting and dual-fiber coupling module

The beam-splitting and dual-fiber coupling module consisted of a laser source and a two-fiber coupling box (**Fig. S2B**). The laser source was a compact, single-wavelength, fiber-based femtosecond system (Alcor920-2W-20MHz, SparkLaser, Martillac, France; 920 ± 10 nm, 100 fs, 20 MHz repetition rate, 2 W output). The laser included integrated group-delay dispersion compensation (0 to −60,000 fs²) and fast analog power modulation (>1 MHz bandwidth). The dispersion compensation range was sufficient to counteract the dispersion accumulated over at least 3 m of the polarization-maintaining hollow-core photonic crystal fiber (HC920-PM, NKT, Copenhagen, Denmark; dispersion at 920 nm: 90 ps/nm/km). After propagating through 2.3 m of HC920-PM, the pulse width (Lorentzian fit) was approximately 100 fs.

Inside the two-fiber coupling box (**Fig. S2C**), the input laser beam was first split into two beams by a polarization beam splitter (PBS122, Thorlabs, NJ, USA; ID #11 in **Fig. S2C** and the **Supplementary Document Part 3: parts list**). The intensity ratio between the two beams was controlled by rotating a half-wave plate (HWP; AHWP05M-980, Thorlabs; ID #10) placed before the PBS. Each beam’s angle and position were independently adjusted using a pair of reflective mirrors (PF05-03-P01-10, Thorlabs; ID #9), mounted in high-stability 2-axis mirror mounts (POLARIS-K05S2, Thorlabs; ID #6). These adjustments were accessible externally without opening the coupling box. Each beam was then coupled into a termination-processed HC920-PM fiber assembly using a coupling lens (C230TMD-B, Thorlabs; ID #12). The coupling procedure followed the protocol described in Zong et al. (Cell, 2022). Briefly, coupling efficiency was monitored by measuring the output power at the distal end of each HC920-PM fiber while iteratively adjusting the mirror pair and the distance between the fiber termination (in its glass flange) and the coupling lens. Coupling efficiencies typically exceeded 80%. Once optimal coupling was achieved, the fiber end, glass flange, and coupling holder were fixed using UV-curable glue (NOA61, Norland, NJ, USA). Due to the minimized number of optical components and the rigid mechanical design, the alignment was highly stable. No active vibration isolation was required, and the output power decreased by <1% over three months, eliminating the need for frequent realignment. An additional HWP was placed in front of each coupling lens to adjust polarization and independently eliminate second-pulse artifacts (Zong et al., Cell 2022). Each beam could be blocked individually using its own shutter (SHB025, Thorlabs; ID #8). In standard dual-region imaging experiments, the first HWP (before the PBS) was adjusted to equalize the power delivered to both miniscopes. When the two imaging regions exhibited different fluorescence intensities, power distribution was fine-tuned by rotating this HWP while monitoring the live image intensities until they were balanced.

Multiple two-fiber coupling boxes can also be used in parallel to drive more than two miniscopes. Commercial multi-beam 2P laser sources provide individual power modulators per output fiber (**Fig. S2G**; ALCOR920-MULTIBEAM, SparkLaser), allowing independent adjustment without affecting other channels. These commercial systems support up to four independent fiber outputs (**Fig. S2G**).

### Dual-miniscope mounting station

The dual miniscope mounting module enabled simultaneous mounting of two miniature microscopes on the animal’s head, each with independent angular and positional adjustment (**Fig. S2D**). The system consisted of two identical miniscope-holding arms. For each arm, we assembled a five-axis motorized scope holder composed of a manual rotation stage (RP03/M, Thorlabs, NJ, USA), an adjustable-angle mounting plate (AP180/M, Thorlabs), and a three-axis piezo-driven linear stage (MMS3, Thorlabs; IDs #41, #42, and #43 in **Fig. S2D** and in the **Supplementary Document Part 3: parts list**). A hardboard plate (TB4, Thorlabs; ID #53) was cut into a 17-cm-diameter circle, coated with rubber spray, and used as a running wheel. Two custom headbar holders (ID #49) were mounted on dual-axis adjustable-angle stages (XRNG-K1/M, Thorlabs; ID #32). Each headbar holder secured and adjusted one implanted headbar from a single side. Together, each miniscope-holding arm combined with its corresponding headbar holder provided full degrees of freedom necessary to align one miniscope precisely to its corresponding cranial window. Importantly, this design allowed independent alignment of each miniscope without needing to compensate for angle differences between the two cranial windows.

### Control and acquisition for the use of two miniscopes

Control and data acquisition for the dual-miniscope setup were performed using a vDAQ system (4-channel, MBF Bioscience, VT, USA) in combination with ScanImage software. A single MEMS driver (PicoAmp 5.4 T180, Mirrorcle, CA, USA) was used to drive both MEMS scanners in the two miniscopes simultaneously. Two TLens drivers (KPC101, Thorlabs, NJ, USA) provided independent high-voltage control of the Tlenses in each miniscope. A high-voltage driver (KPC101, Thorlabs, NJ, USA) provided bias voltage to all SIPMS in two miniscopes. Two piezo-driven three-axis micromanipulators (uMp-TSC, Sensapex, Oulu, Finland) were used to position and baseplate each miniscope independently, controlled directly through ScanImage. The photocurrent outputs from the four SiPM detectors (two per miniscope) were routed through four transimpedance amplifiers (DHPCA-100_FEMTO, FEMTO, Berlin, Germany). The amplified analog signals were then digitized by the four high-speed analog-to-digital converter channels of the vDAQ (125 Mbit/s per channel).

### Electronic cable assembly

The electronic cable assembly (**Fig. S2E**) delivered control signals to the MEMS scanners and Tlenses of the two MINI2P-II miniscopes and transmitted the photocurrent signals from the SiPMs to the remote transimpedance amplifiers. The assembly consisted of 8 × 36-AWG silver-plated copper wires (Ø = 0.3 mm; Microwire, GA, USA) and 4 × micro-coaxial cables (PCX36K10AK, AXON’ CABLE, France). Six of the silver-plated copper wires were split in the final 10 cm of the assembly, providing identical signals to both miniscopes. These shared lines carried the 2-axis MEMS scan signals (wire IDs #1, #2, #3, and #4), a common ground (wire ID #5), and the SiPM bias voltage (0 to +60 V). The remaining two silver-plated wires delivered independent high-voltage control signals for the Tlenses in each miniscope. Each SiPM transmitted its photocurrent through a dedicated micro-coaxial cable. Within each miniscope, the MEMS and Tlens components were soldered onto a single flexible PCB (ID #i in the **Supplementary Document Part 3: parts list**; PCB design available via GitHub upon acceptance of the manuscript or can be provided upon request.) that interfaced with the main cable via a compact 6-pin micro-connector. The two SiPMs within each miniscope were connected to a second flexible PCB (ID #ii in the shopping list; PCB design also available via GitHub or can be provided upon request.). This lightweight, compact cabling architecture minimized mechanical load and torsion on the animal while maintaining robust signal integrity and reliable control of all active microscope components.

### Evaluating polarization-maintaining property of HC920 vs HC920-PM

To compare the polarization-maintaining properties of HC920 versus HC920-PM, both fibers were integrated into a dual-fiber laser coupling module using a beam splitter to ensure reliable, side-by-side comparison. For each measurement, one fiber at a time was connected to a polarimeter (PAX1000IR1/M, Thorlabs). The tested fiber was laid flat and straight on a leveled surface to minimize unintentional stress. During bending measurements, the fiber was wrapped around custom made 3D-printed disks of defined radians. After adjusting the fiber onto the disk, we waited 0.5-1 minute for the output polarization from the fiber to be stabilized before recording. This procedure was repeated independently for each bending radius. For twist tests, the fiber was attached to a disk of 80-mm diameter, chosen to impose minimal mechanical constraint on the fiber. The disk was rotated in 360° increments to introduce controlled amounts of twist. After each additional turn, the fiber was again allowed to stabilize for 0.5–1 min before measurement. Each measurement consisted of a constant amount of 1000 data points.

## QUANTIFICATION AND STATISTICAL ANALYSIS

### 2P imaging data preprocessing

a. Dataset preparation

For single-scope imaging (data in **Fig.1** and **Fig. 6**), preprocessing of 2P time-series stacks followed the procedure described previously (Zong2022). For dual-scope imaging, raw recordings were first split into two datasets using a custom-made MATLAB script. Each dataset recorded from one miniscope was then processed independently using the standard preprocessing pipeline described briefly below.

1. b. Distortion correction.

Although MINI2P-II eliminates much of the scanning field bending and magnification variation caused by the MEMS mounting angle in MINI2P, a small residual geometric distortion remains due to MEMS scanner nonlinearity (**Fig.1E-i**). We corrected this distortion using the approach described in Zong et al., 2022. Briefly, we generated a set of transformation matrices for different zoom levels and Tlens focus positions, mapping the raw MINI2P-II image to an ideal undistorted target. These matrices were generated using images of a fluorescent gird sample (50 µm grid, R1L3S3P, Thorlabs). For each zoom and Tlens position within 150 µm *z*-range, we first imaged the grid repeatedly, then identified anchor points were at each grid intersection, and computed a piecewise-linear 2D transformation was in MATLAB, relating distorted positions to true coordinates. We applied this matrix to each imaging data before downstream analysis in Suite2p.

1. c. Motion correction

Motion artifacts in awake animals arise from breathing, head movement, and brain swelling and -shrinkage introduced during imaging. We used *Suite2P* to conduct a two-step motion correction which first performs rigid registration (assuming that the whole frame of image shifts by the same amount of distance), and then non-rigid registration (assuming that subsegments of each frame of image shift by separate amounts). Rigid registration computes the global shifts *dx* along the *x*-axis and *dy* along the *y*-axis between each frame and the reference image using phase-correlation. Non-rigid registration divides the image into blocks with a certain fraction of overlap. In our recordings, each block has 64×64 pixels and the whole FOV was partitioned into 12×12 (for 512 × 512 image) or 6×6 (for 256 × 256 image) partially overlapping blocks. The shift of each block is calculated separately. Then the shift in each block is smoothed using bilinear interpolation in 2D to create the final shift matrixes on the *x*-axis, and *y*-axis. Throughout the manuscript, rigid motion is reported as the per-frame d*x* and d*y* displacements between each frame and the reference frame.

1. d. Neuron detection and signal extraction

After motion correction, *Suite2p* was used to detect neurons, extract fluorescence traces and perform calcium-trace deconvolution. Deconvolution used a non-negative algorithm with an exponential kernel (decay τ = 1.5 s for GCaMP6s), consistent with previously published parameters (Friedrich et al., 2017; Pachitariu et al., 2017). Deconvolution increases temporal precision and denoises fluorescence signals. Suite2p outputs (raw fluorescence, neuropil signal, and deconvolved activity) were processed using NATEX, a custom pipeline for neuronal activity processing, behavior-tracking processing and alignment of activity with behavioral time-series.

Next, the *Suite2p* output, including the fluorescent, neuropile and the deconvolved signals extracted for each neuron, is fed into software NATEX for processing the neural activity and tracking data. It includes three blocks, related to neuronal activity processing, processing of tracking data, and alignment and combination of neuronal activity and tracking data. We used NATEX to calculate the fractional change in fluorescence with respect to baseline (ΔF/F) of each neuron and detect significant transients which were defined as transients that occur in periods in which ΔF/F exceeds 2 times the local standard deviation of the baseline fluorescence for more than 0.75s (Zong et al., 2022). Deconvolved traces within significant-transient windows were rescaled so that their maximum matched the ΔF/F maximum. Non-zero deconvolved events are referred to as “calcium events.” Spatial tuning analysis in RSC/PPC (**Figs. 5, S5**) used deconvolved calcium events and all other analyses used ΔF/F.

1. e. SNR calculation.

For each neuron, signal-to-noise ratio (SNR= signal / noise) was calculated as i) signal: the mean amplitude of all 90th-percentile ΔF/F values within significant-transient windows; ii) noise: the mean of the frame-to-frame ΔF/F differences outside significant-transient windows.

### Tracking data processing

The position of mice during experiments was analyzed in DeepLabCut (DLC) (Mathis et al., 2018) from raw video recordings. Each video frame resulted in estimated body parts (miniscope, left ear, right ear, body center, and tail base) position coordinates.

The DLC model for tracking the horizontal position of the animal in 80x80 cm recording box was trained using 11 video recordings from different mice obtained on different experiment days. 220 frames were extracted from these videos (20 frames per video) and manually labeled. The DLC output provided, for each frame, the pixel coordinates of the tracked body parts along with a confidence score vector (ranging from 0 to 1) that reflects the likelihood of each position estimate. The frames were filtered using a built-in DLC median filter when the likelihood values did not reach the threshold of 0.5. The head location was computed as the midpoint between the left and right ear positions and represented by the x–y coordinates *X*_ℎ*ead*_(*t*) and *Y*_ℎ*ead*_(*t*). The traces Xhead(t) and Yhead(t) were smoothed by computing a linear regression in each window of 0.2 s (3 frames) independently.

For ladder climbing behavior, two DLC models were combined, one tracking the horizontal position (DLC trained on 13 recordings with 260 labeled frames in total, same as before), the other considering the vertical position of the mice (28 recordings, with 16 frames extracted from each considering only miniscope position). Climbing periods were manually noted based on the raw video recordings and relevant behavior windows were selected from DLC output coordinates.

### Rastermap analysis

To visualize population activity, neurons were sorted using Rastermap algorithm (Stringer et al., 2025). Rastermap was applied separately to left and right hemisphere neurons using the following updated parameters: number of clusters=64, number of principal components=32, locality=0.1, time lag window=15, grid upsample=0. The resulting neuron ordering was used to plot activity raster. Population activity for each hemisphere was computed as the sum of ΔF/F across neurons at each timepoint. The summed signal was z-scored across time, smoothed using a Gaussian filter (σ = 5 frames), and shifted so that the minimum value was zero for visualization.

### Principal Component Analysis (PCA)

We compared population dynamics across four sampling conditions: (1) left hemisphere only, (2) right hemisphere only, (3) all left and right hemisphere neurons, and (4) a balanced left–right subsample in which neurons were drawn equally from both hemispheres but matched in total count to the unilateral conditions. Neural activity matrices (neurons × time) were first z-scored and temporally smoothed using a Gaussian filter (σ = 15 frames) to reduce high-frequency noise while preserving task-related structure. Behavioral data were segmented into climbing-up and climbing-down epochs. Each epoch was temporally normalized to a fixed length (n=100 samples) using linear interpolation, enabling comparison across variable-duration episodes. For each condition, concatenated epoch data were used to compute a shared principal component basis using PCA. To control sampling variability, this procedure was repeated 100 times, each time subsampling 150 neurons per condition. Each epoch was then projected into the resulting neural manifold, and trajectories were visualized in the first three principal components.

### Subspace angle calculation

To quantify the similarity between climbing-up and climbing-down population dynamics, we compared their low-dimensional subspaces using principal angle analysis. For each condition and epoch, population activity was reduced via PCA, retaining the top *k* = 15 components. This yielded orthonormal basis matrices *U* ∈ ℝ*^n^*^×*k*^ and *V* ∈ ℝ*^n^*^×*k*^ corresponding to the climbing-up and climbing-down conditions, respectively. Subspace similarity was assessed by computing the singular value decomposition (SVD) of the cross-subspace alignment matrix: *M* = *U*^T^*V*. The singular values *σ_i_* of *M* correspond to the cosines of the principal angles *θ_i_* between the two subspaces, i.e., *σ_i_*= cos(*θ_i_*). We defined the mean subspace angle as 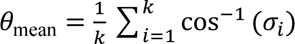 which captures the average angular separation between the climbing-up and climbing-down trajectories. This metric was computed independently for each epoch and subsequently aggregated across conditions. For visualization, all angles were expressed in degrees.

### Allocentric spatial tuning maps

Allocentric spatial tuning maps were generated by normalizing each neuron’s calcium activity by the animal’s occupancy of each spatial bin. For each bin, the tuning value was computed as the ratio between (1) the sum of the amplitudes of the neuron’s deconvolved calcium activity occurring within the bin and (2) the total time the animal spent in that bin. Spatial bin sizes were 2.5 × 2.5 cm². All maps were smoothed with a Gaussian kernel (standard deviation: 3 cm). The color scale in spatial heatmaps ranged from zero to the maximal bin value within the session, expressed in ΔF/F·s⁻¹. To rule out effects of changes in behavioral state (foraging versus sitting still), we dismissed all data at running speeds lower than 2.5 cm/s in all calculations of the observed and shuffled data.

### Egocentric boundary rate maps (EBRs)

Egocentric boundary rate maps (EBRs) were generated using a modified version of the method described in Alexender, et.al, 2020. For each video frame, the positions of the arena boundaries (i.e., the box walls) were computed relative to the animal’s instantaneous position. At every time point, we measured the distance between the mouse and the surrounding walls in 2.5-cm radial bins, and the angular position of the walls in 3° angular bins spanning 0–360°. Angular bins were aligned to the animal’s head direction, such that: 0° corresponded to directly in front of the mouse, 90° to its left, 180° directly behind, and 270° to its right. For each bin (2.5 cm × 3°), the EBR value was computed as the ratio of the sum of the amplitudes of the deconvolved calcium activity, and the total occupancy time within that bin across the full recording session. The resulting rate maps were smoothed with a Gaussian kernel (standard deviation: 5 cm × 6°).

### EBCs in RSC and PPC

For each neuron with more than 100 calcium events in a given session, we quantified its egocentric boundary tuning using standard circular statistics as described previously (Alexender, et.al, 2020). Specifically, we computed the mean resultant vector (MR), reflecting the neuron’s egocentric boundary directional firing after collapsing across the distance dimension; the mean resultant length (MRL), defined as the absolute magnitude of MR and representing the strength of egocentric bearing tuning; and the mean resultant angle (MRA), indicating the neuron’s preferred egocentric boundary orientation. MRL was computed for each neuron over the entire 40-min session, whereas MRA and preferred distance were calculated separately for the first and second 20-min halves to evaluate tuning stability. A neuron was classified as an egocentric boundary cell (EBC) if it met all of the following criteria, identical to those used in Alexender, et.al, 2020, 1) the neuron’s baseline MRL exceeded the 99th percentile of the MRL distribution from shuffled data; 2) the absolute circular difference in MRA between the two session halves was < 45° and 3) the change in preferred distance for each half relative to the full session value was < 50%.

Shuffling was performed independently for each neuron using 200 permutations. For each permutation, the full sequence of the neuron’s deconvolved calcium events was time-shifted along the animal’s trajectory by a random interval selected uniformly between +30 s and (session length − 30 s). Session boundaries were treated cyclically such that data wrapping from the end to the beginning was seamless.

### Statistical tests

All statistical tests were two-sided, non-parametric tests because the distributions of statistics were not assumed to be normally distributed. Statistical significance was defined as p < 0.05. We used Shapiro–Wilk test for normality and paired t-tests or Wilcoxon signed-rank tests as appropriate to assess the influence of miniscope weight and cable flexibility on animal behavior. A Mann– Whitney U test was used to compare imaging quality between left and right M1 hemispheres. To test whether mean ΔF/F differed between left and right M1 within each behavioral state, we used the Wilcoxon signed-rank test (two-sided), treating mean ΔF/F from each hemisphere as paired observations. To test whether mean ΔF/F differed across the seven behavioral states within each hemisphere, we used the Kruskal–Wallis test. To test whether bilateral correlation differed between specific pairs of behavioral states, we used the Wilcoxon rank-sum test (two-sided). To evaluate the predominance of leftward versus rightward (contralateral) receptive fields in RSC/PPC EBCs, we applied a binomial test and a Rayleigh test, the latter being appropriate for identifying non-uniformity or periodicity in circularly distributed data. To test whether the two full circular distributions of preferred orientations differ from each other, we use Watson’s two-sample U² test — the circular analogue of the two-sample KS test.

### Additional resources

Updated versions of all key component ordering numbers, 3D models and engineering drawings for custom parts, detailed assembly instructions, video tutorials, and customized recording software will be made available on GitHub upon acceptance of the manuscript or can be provided upon request.

## Notes

### Summary of Updates

The previous author list was incorrect, which was corrected in the new version.

